# Abnormal antibodies to self-carbohydrates in SARS-CoV-2 infected patients

**DOI:** 10.1101/2020.10.15.341479

**Authors:** Dorothy L. Butler, Jeffrey C. Gildersleeve

## Abstract

SARS-CoV-2 is a deadly virus that is causing the global pandemic coronavirus disease 2019 (COVID-19). Our immune system plays a critical role in preventing, clearing, and treating the virus, but aberrant immune responses can contribute to deleterious symptoms and mortality. Many aspects of immune responses to SARS-CoV-2 are being investigated, but little is known about immune responses to carbohydrates. Since the surface of the virus is heavily glycosylated, pre-existing antibodies to glycans could potentially recognize the virus and influence disease progression. Furthermore, antibody responses to carbohydrates could be induced, affecting disease severity and clinical outcome. In this study, we used a carbohydrate antigen microarray with over 800 individual components to profile serum anti-glycan antibodies in COVID-19 patients and healthy control subjects. In COVID-19 patients, we observed abnormally high IgG and IgM antibodies to numerous self-glycans, including gangliosides, *N*-linked glycans, LacNAc-containing glycans, blood group H, and sialyl Lewis X. Some of these anti-glycan antibodies are known to play roles in autoimmune diseases and neurological disorders, which may help explain some of the unusual and prolonged symptoms observed in COVID-19 patients. The detection of antibodies to self-glycans has important implications for using convalescent serum to treat patients, developing safe and effective SARS-CoV-2 vaccines, and understanding the risks of infection. In addition, this study provides new insight into the immune responses to SARS-CoV-2 and illustrates the importance of including host and viral carbohydrate antigens when studying immune responses to viruses.

## Introduction

COVID-19 is a respiratory disease caused by the severe acute respiratory syndrome coronavirus 2 (SARS-CoV-2). In less than a year, this virus has caused over 1 million deaths worldwide and has become the third leading cause of death in the United States.^1^ Beyond the severe impact on human health, SARS-CoV-2 has caused major disruptions to many aspects of life, including the economy, education, travel, and personal life. As a result, an unprecedented global effort is underway to develop effective methods to prevent and treat COVID-19. Because this is a new, emerging infectious virus, much of the fundamental knowledge that provides the foundation for developing vaccines and therapeutic agents, as well as for making informed public health decisions, is lacking. Therefore, there is an urgent need to improve our basic understanding of how the virus works, why it causes severe disease outcomes, and how we can intervene to protect human life.

One of the most perplexing aspects of the disease is that it can cause a myriad of symptoms in addition to respiratory distress, often involving multiple organs apart from the lungs. For example, COVID-19 patients can suffer from a range of neurological symptoms, including encephalopathy, psychosis, neurocognitive syndrome, and headaches.^2–5^ Beyond impacting neurological functions, SARS-CoV-2 infections have also been reported to affect the cardiovascular and gastrointestinal systems.^6–9^ An especially troubling issue is that some symptoms can last for months beyond the primary infection, even in the absence of detectable virus.^10–12^ It is unclear why some patients, often referred to as “long haulers,” have prolonged effects. More generally, the specific mechanisms that lead to disparate symptoms and damage in multiple organs are not well understood.

Our immune system plays a critical role in preventing, clearing, and treating SARS-CoV-2. Therefore, understanding host immune responses to SARS-CoV-2 is essential for developing effective therapies and vaccines to control this pandemic. While the immune response can involve many elements of the innate and adaptive arms of the immune system, antibody responses are one of the most important features. Most patients develop a robust antibody response to the virus, and the presence of these antibodies can be used as an indicator of recent infection.^13–15^ The presence of neutralizing antibodies in recovering patients has also been exploited for treating new infections through the administration of convalescent serum.^16–19^ Neutralizing monoclonal antibodies isolated from patients or identified via *in vitro* techniques are currently in clinical trials for treating COVID-19.^20–22^ Furthermore, the generation of a vigorous antibody response is a key objective for the development of an efficacious vaccine. For these reasons, a thorough understanding of antibody responses to SARS-CoV-2, as well as to vaccines, is vital to these objectives.

While often beneficial, overly aggressive and/or aberrant immune responses can also be harmful in COVID-19 patients.^23–25^ For example, excessive inflammation has been associated with severe respiratory effects and detrimental symptoms.^26, 27^ Awareness of this issue has led to the use of anti-inflammatory agents, such as dexamethasone, to significantly reduce mortality in COVID-19 patients.^28^ Emerging evidence indicates that SARS-CoV-2 may also induce autoantibodies. For example, autoantibodies have been identified in children who have previously had COVID-19 and have relapsed with multi inflammatory syndrome.^29^ Other studies have shown autoantibodies to a variety of proteins in adult patients with severe COVID-19 symptoms or neurological symptoms.^30–32^ Autoantibodies to certain gangliosides have also been observed in a subset of COVID-19 patients with Guillain-Barre Syndrome (GBS) related symptoms.^33, 34^ Lastly, certain antibody responses can actually enhance infection,^35^ but the mechanisms of antibody-dependent enhancement are not well understood. For these reasons, studying the host immune response, especially the antibody response, is also critical for understanding complications that can arise from an overly aggressive immune response and for developing interventions to circumvent these problems.

Numerous groups have been studying immune responses to SARS-CoV-2, and a wealth of new information is emerging.^23–25, 29, 30, 32, 36–44^ Although roles of various cells, cytokines, and antibodies to proteins are being uncovered, relatively little is known about immune responses to carbohydrates. Some recent reports have shown a small correlation with ABO blood type and susceptibility to COVID-19, and this effect may involve pre-existing serum antibodies to the blood group A (BG-A) and/or blood group B (BG-B) carbohydrates.^36–39^ Another recent study reported an inverse relationship between COVID-19 disease severity and serum anti-α-Gal antibodies.^45^ α-Gal is a non-human glycan, and natural antibodies to this glycan epitope can be part of the protective response to pathogenic viruses, bacteria, and parasites that contain this glycan.^45–48^ In addition to these studies on serum anti-carbohydrate antibodies, several studies have demonstrated that the SARS-CoV-2 spike protein is heavily glycosylated.^49–54^ Glycosylation mapping of the spike protein subunits revealed a variety of *O*-linked and *N*-linked glycans, including high-mannose.^50^ These glycans can be recognized by 2G12, an antibody that targets high mannose glycans on gp120 of HIV.^55^ Collectively, these studies suggest that glycans and anti-glycan antibodies may play an important role in the prevention, treatment, and severity of COVID-19.

To better understand the roles of glycans in the immune response to SARS-CoV-2, we compared serum anti-glycan IgG and IgM antibody repertoires of 40 COVID-19 patients with 20 uninfected control subjects. To monitor a large and diverse assortment of antibody populations, we profiled each serum sample using a carbohydrate antigen microarray with over 800 components. These studies revealed that COVID-19 patients had substantial differences in anti-glycan antibodies, including unusual antibodies to a variety of self-glycans.

## Results

### Study design

Serum from 40 SARS-CoV-2 infected patients and 20 uninfected individuals were used in the study. All control serum samples were collected before December 2019 when the outbreak of SARS-CoV-2 began. There were 20 male and 20 female individuals in the COVID-19 cohort, and there were 13 male and 7 female individuals in the control group. All patients in the COVID-19 cohort had a positive antibody test for IgG, IgM, or both to the spike protein receptor binding domain using an indirect ELISA. All patients were symptomatic, but details about specific symptoms and outcomes were not available at the time of this study. The average patient age of those infected with SARS-CoV-2 was 64, with an age range of 41-92 years old. The uninfected, control individuals had an average age of 40, with an age range of 18-65. This difference in age between the control group and the COVID-19 positive group may have some influence on the results (see below).

To assess the anti-glycan repertoires of patients with COVID-19, we profiled IgG and IgM from serum samples on a carbohydrate antigen microarray containing 816 components. The microarray included a diverse collection of *N*- and *O*-linked glycans, glycolipid glycans, glycopeptides, bacterial and fungal glycans, and some natural glycoproteins. This set of glycans allows for rapid profiling of a broad range of anti-glycan antibody populations in serum including those to both foreign and self-antigens. Antibody signals from each COVID-19 patient were compared to the control set to identify unusual signals.

### Overall profiles reveal significantly lower IgM signals in in SARS-CoV-2 positive patients

We started by evaluating overall antibody signals across the array to assess global differences in antibody levels in control and COVID-19 patient samples and to provide context for individual differences. We measured the mean IgG and IgM signals from all the array components for each cohort of samples (Figure 1). For nearly every glycan except a few detailed below, the mean IgM signals to glycans were 2 to 4 fold lower in SARS-CoV-2 positive patients compared to controls, while the total mean IgG signals were similar. Across the entire array, the average IgM signals in the control group were 2.3-fold higher than COVID-19 patients. To determine if this effect was specific to carbohydrate-binding IgM or due to differences in total serum IgM levels, we measured the total IgM in all samples. The average total IgM in the COVID-19 patient samples was 30% lower than the average total IgM in the control samples (Supplemental Figure S1). Thus, differences in total IgM only partially explain the substantially lower IgM signals observed on the array in SARS-CoV-2 positive patients. We have previously observed large decreases in carbohydrate-binding IgM with increasing age.^56^ Therefore, differences in the average ages of each sample population are likely to also contribute.

**Figure 1:**
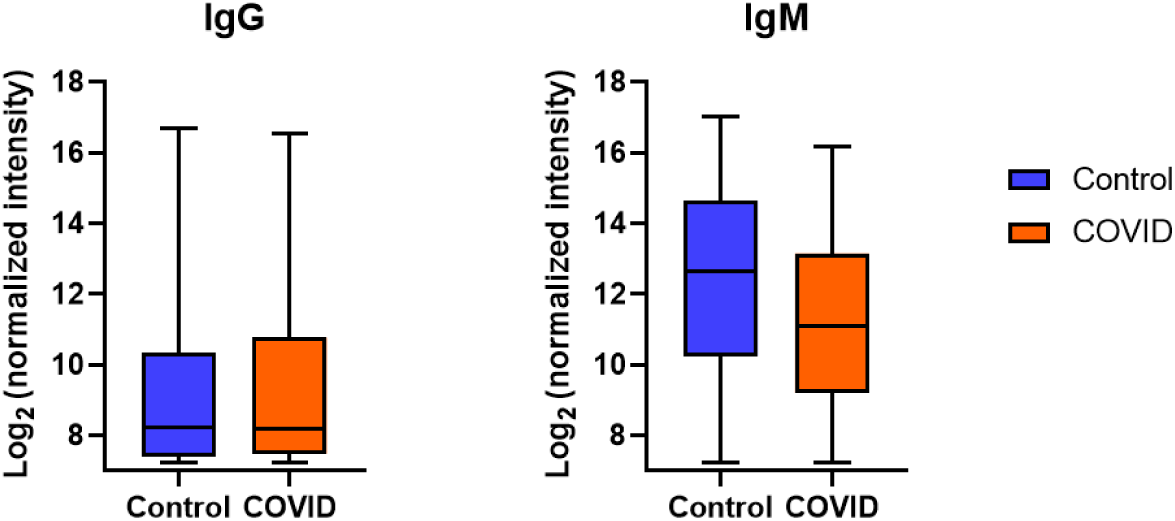
Average IgG and IgM antibody signals to all glycans. Box and whisker plots of the average signals (log-transformed base 2) to all array components for IgG and IgM antibodies from control and COVID-19 serum samples.

### Unusually high IgG to glycolipids in SARS-CoV-2 positive patients

One striking difference between serum samples from COVID-19 patients and healthy controls were unusually high antibodies to glycolipid glycans (see Figure 2 and Supplemental Figure S2). Unusually high was defined as a signal that was greater than 6 standard deviations above the mean of the control group and greater than 10-fold above the floor value for our assay. While very uncommon in healthy individuals, anti-glycolipid antibodies are often found in populations that have autoimmune diseases and other nervous system dysfunctions.^57^ For example, antibodies to asialo-GM1, GM1a, GD1a, and GD1b are frequently observed in patients with Guillain-Barre Syndrome (GBS). We observed unusually high antibodies to GBS glycans in 15% of patients (Figure 2A). Even larger signals were observed to several other glycolipids not associated with GBS, such as GD3, fucosyl-GM1, GM2, and GM3 (see Figure 2B). The largest antibody signals for GD3 and fucosyl-GM1 in COVID-19 patients were >35-fold higher than the largest signals in the control group. Although humans do not biosynthesize Neu5Gc, it can be obtained via dietary sources and incorporated into cell surface glycans;^58^ therefore, we have included the Neu5Gc variant of GD2 [#505; GD2 (Gc/Gc)] with this group (see Figure 2B). When considering all the glycolipids (GBS and non-GBS), 14 patients (35%) had high antibodies to at least one glycolipid. Antibody signals to gangliosides were not correlated with IgG titers to the spike protein.

**Figure 2.**
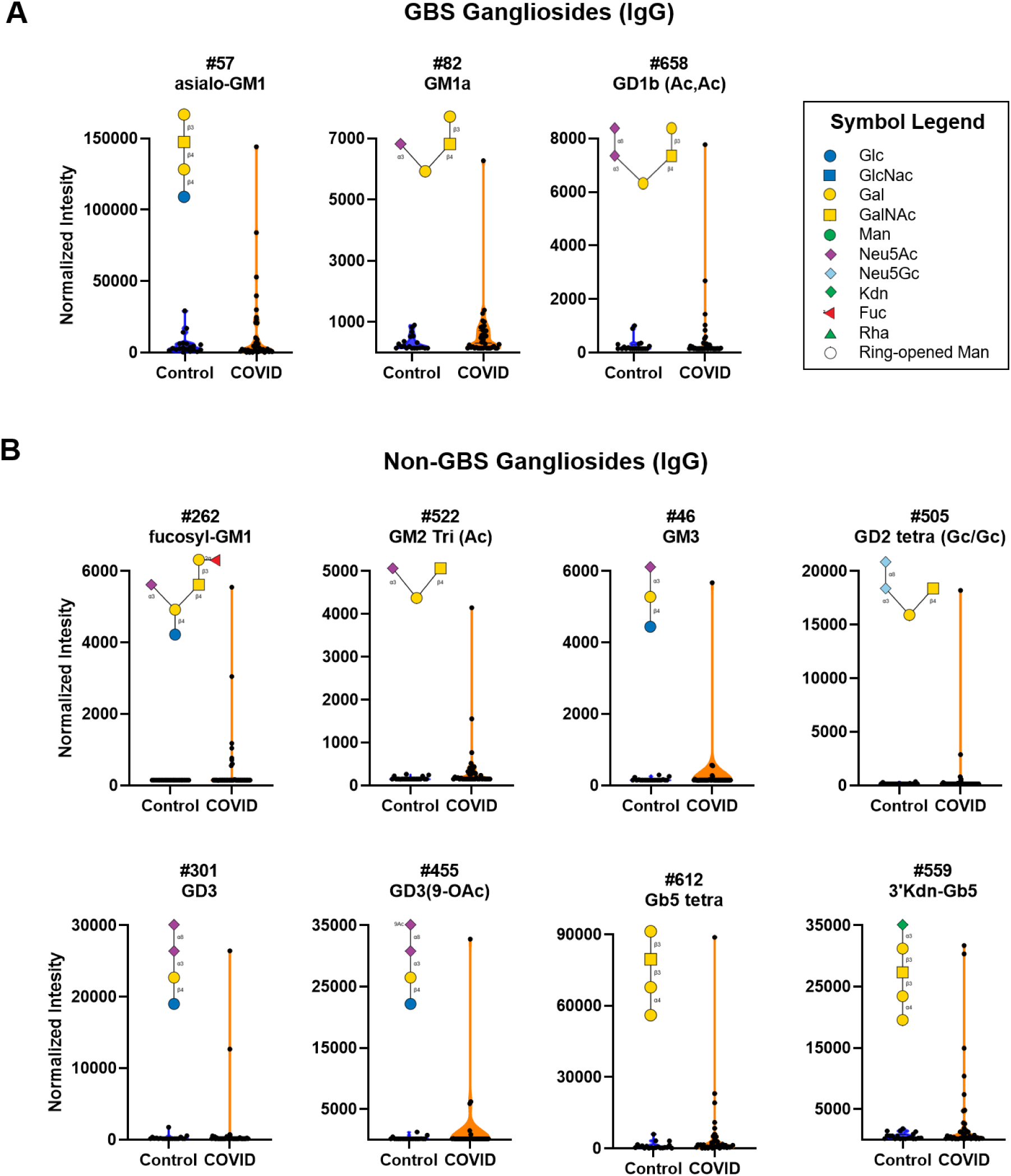
High antibody signals to select ganglioside glycans in COVID-19 patient serum. Violin plots showing high IgG signals to various gangliosides/glycolipids for COVID-19 patients versus control subjects, with each point representing data from an individual patient: **A)** Guillain-Barre Syndrome (GBS)-associated ganglioside and **B)** other gangliosides/glycolipids. See Figure 6 for patients with signals to multiple glycans. Glycan structures were created using GlycoGlyph.^59^

### Unusually high serum antibody signals to oligomannose and other N-linked glycans in SARS-CoV-2 positive patients

In addition to the antibodies to glycolipids, we also observed unusually large IgG signals to *N*-linked glycans and oligomannose fragments of certain *N*-linked glycans (see Figure 3, Figure 4, Supplemental Figure S3 and S4). *N*-linked glycans are abundant in the human body, and they also cover the spike proteins of SARS-CoV-2. Our array contains approximately 30 different *N*-linked glycans, including high mannose, complex, and hybrid *N*-glycans. Overall, there were very little or no measurable signals for *N*-linked glycans in the control group. In contrast, there were a variety of noticeably high serum antibody signals to several *N*-linked glycan in the SARS-CoV-2 positive patients (see Figure 3 and Figure S3). The largest and most unusual IgG signals were to NGA4, a complex, tetraantennary *N*-glycan with the following sequence: GlcNAcβ1-2(GlcNAcβ1-6)Manα1-6[GlcNAcβ1-2(GlcNAcβ1-4)Manα1-3]Manβ1-4GlcNAcβ. Four patients had high antibodies to NGA4, and the largest signal to NGA4 in the COVID-19 group was greater than 40-fold higher than the largest signal in the control group. Interestingly, only one patient had high antibody signals to the corresponding triantennary *N*-glycan, NGA3 (GlcNAcβ1-2Manα1-6[GlcNAcβ1-2(GlcNAcβ1-4)Manα1-3]Manβ1-4GlcNAc), and none had high signals for the biantennary *N*-glycan NGA2 (GlcNAcβ1-2Manα1-6(GlcNAcβ1-2Manα1-3)Manβ1-4GlcNAc). Moreover, high IgG signals were not observed for NGA3B (GlcNAcβ1-2Manα1-6[GlcNAcβ1-2(GlcNAcβ1-4)Manα1-3](GlcNAcβ1-4)Manβ1-4GlcNAc). These results indicate a specific response to NGA4 and NGA3, rather than a non-specific or polyreactive response. In addition to NGA4, multiple patients had high IgG signals to Man6-I, a high mannose *N*-glycan with the sequence: Manα1-6(Manα1-3)Manα1-6(Manα1-2Manα1-3)Man. Also of note, one patient showed an unusually high IgG signal to two biantennary sialylated *N*-glycans: “A2 (a2-3)” with the sequence Neu5Acα2-3Galβ1-4GlcNAcβ1-2Manα1-6(Neu5Acα2-3Galβ1-4GlcNAcβ1-2Manα1-3)Manβ1-4GlcNAcβ, and “A2 (a2-6)” with the sequence Neu5Acα2-6Galβ1-4GlcNAcβ1-2Manα1-6(Neu5Acα2-6Galβ1-4GlcNAcβ1-2Manα1-3)Manβ1-4GlcNAcβ. Overall, 17.5% of COVID-19 patients had high IgG signals to 1 or more *N*-linked glycans and 10% had high IgG signals to 2 or more *N*-linked glycans. Abnormally high signals to *N*-linked glycans were also observed for IgM, such as antibodies to A2 (a2-6), Man6-I, and Man9 (see Figure 3B). Most patients with high IgM to *N*-glycans were distinct from patients that had high IgG to *N*-glycans; only three patients had both high IgG and high IgM to *N*-glycans. High antibody signals to *N*-glycans are especially remarkable given that total IgM and IgM signals to the vast majority of other glycans were lower for COVID-19 patients. Neither IgG nor IgM signals to *N*-linked glycans were correlated with titers to the spike protein.

**Figure 3.**
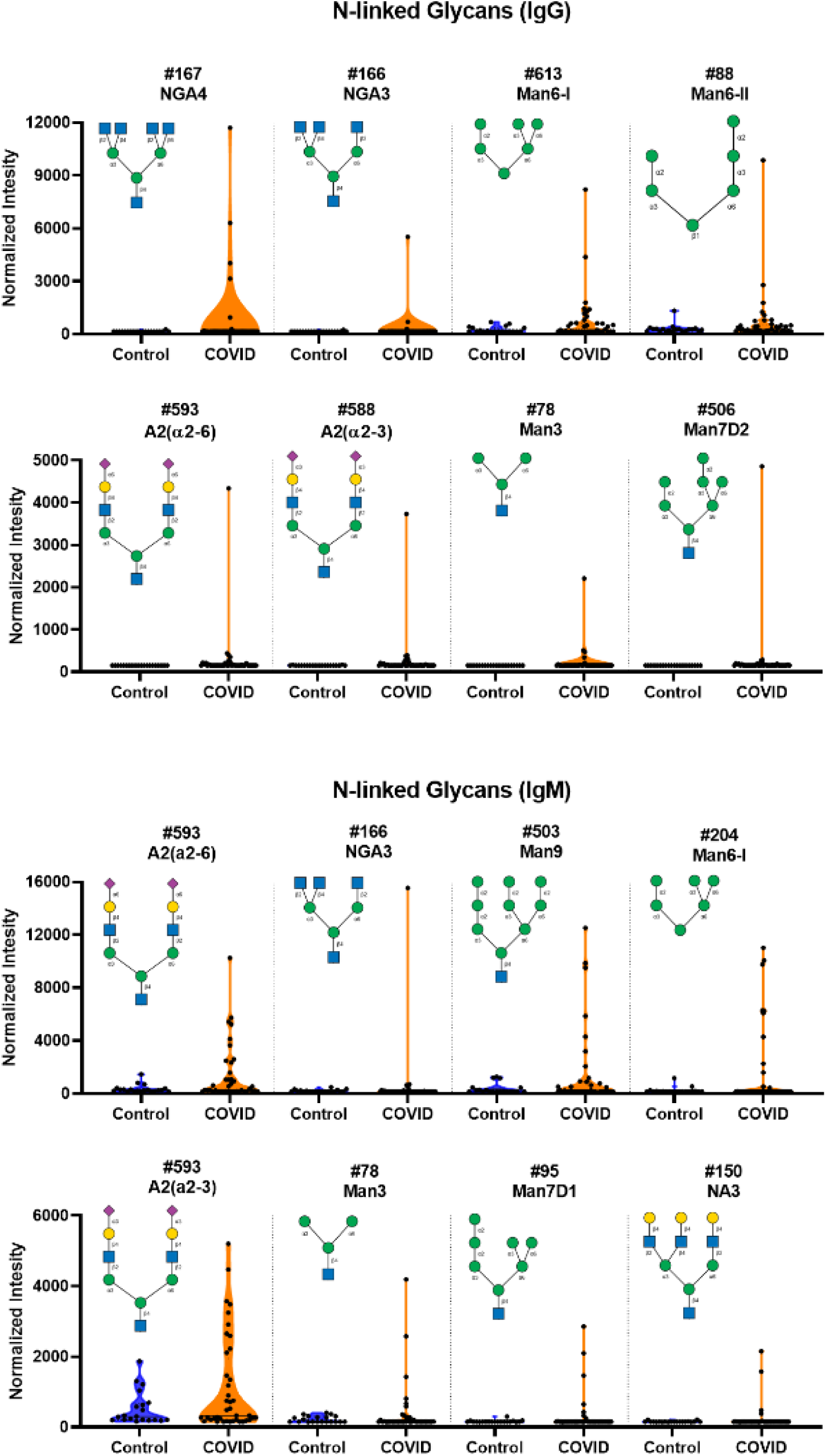
High IgG and IgM signals to select N-linked glycans in COVID-19 patient serum. **A)** Violin plots show several high IgG signals to select *N*-linked glycan array components for serum from COVID-19 patients compared to baseline signals seen from serum from control donors, with each point representing data from an individual patient. **B)** Violin plots show several high IgM signals to select N-linked glycan array components for serum from COVID-19 patients compared to baseline signals seen from serum from control donors. See Symbol Key in Figure 1. See Figure 6 for patients with signals to multiple glycans. Glycan structures were created using GlycoGlyph.^59^

**Figure 4.**
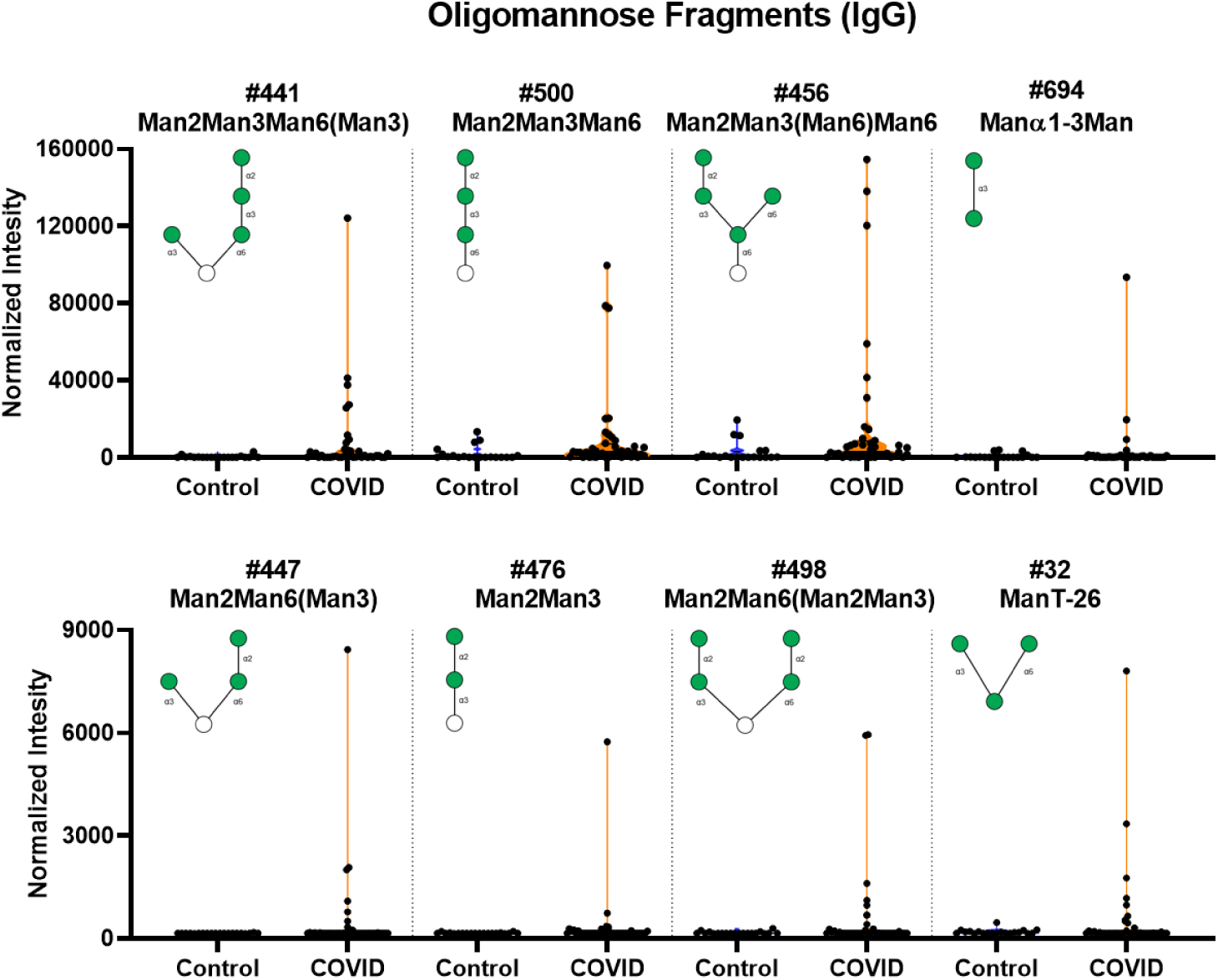
High IgG signals to select oligomannose fragments in COVID-19 patient serum. Violin plots show several high IgG signals to select oligomannose glycan array components for serum from COVID-19 patients compared to baseline signals seen from serum from control donors, with each point representing data from an individual patient. See Symbol Key in Figure 1. See Figure 6 for patients with signals to multiple glycans. Glycan structures were created using GlycoGlyph.^59^

We also observed abnormally high IgG signals for a variety of oligomannose glycans (see Figure 4 and Figure S4). These glycans are substructures or fragments of various *N*-linked glycans. IgG signals for these glycans are typically low in healthy subjects and were low in our control group. Certain patients, however, had very high signals to these glycans. The largest signals were to oligomannose glycans containing a Manα1-2Manα1-3Manα1-6 sequence, but high signals were also observed to several other variants. Overall, 52.5% of COVID-19 patients had high signals to 1 or more oligomannose fragments, and 37.5% had high signals to 2 or more oligommanose fragments. There was a positive association of higher IgG signal for oligomannose fragments with age. No correlation was observed with IgG titers to the spike protein. For IgM antibody signals, there were only small differences for oligomannose fragments.

### Unusually high serum IgM antibody signals to LacNAc and other self-glycans in SARS-CoV-2 positive patients

One common carbohydrate structure found on many *N*-linked glycans, *O*-linked glycans, and glycolipids is N-acetyllactosamine (LacNAc; Galβ1-4GlcNAc).^60^ LacNAc is abundant in humans and many other organisms and can be present as a single unit, as an oligomer of several units, or as longer poly LacNAc repeats. Because LacNAc is abundant in humans, it is considered a “self” glycan. Several COVID-19 patients displayed markedly high IgM signals to LNnO, a glycan containing 3 LacNAc units attached to a galactose residue (Galβ1-4GlcNAcβ1-3Galβ1-4GlcNAcβ1-3Galβ1-4GlcNAcβ1-3Gal). High IgM to this glycan in COVID-19 patients is especially notable given that total IgM and IgM to most other glycans were much lower in COVID-19 patients. In addition, one patient also had a large IgG signal to this glycan (Figure 5). This signal was over 150 fold higher than the largest signal to LNnO in the control group. Little or no measurable signals were detected in our control group for either IgG or IgM. Some differences were also detected for LacNAc and LNnT (Galβ1-4GlcNAcβ1-3Gal) (see Figure 5A and Supplemental Figure S5A). Antibody signals to LacNAc derivatives were not correlated with titers to the spike protein.

**Figure 5.**
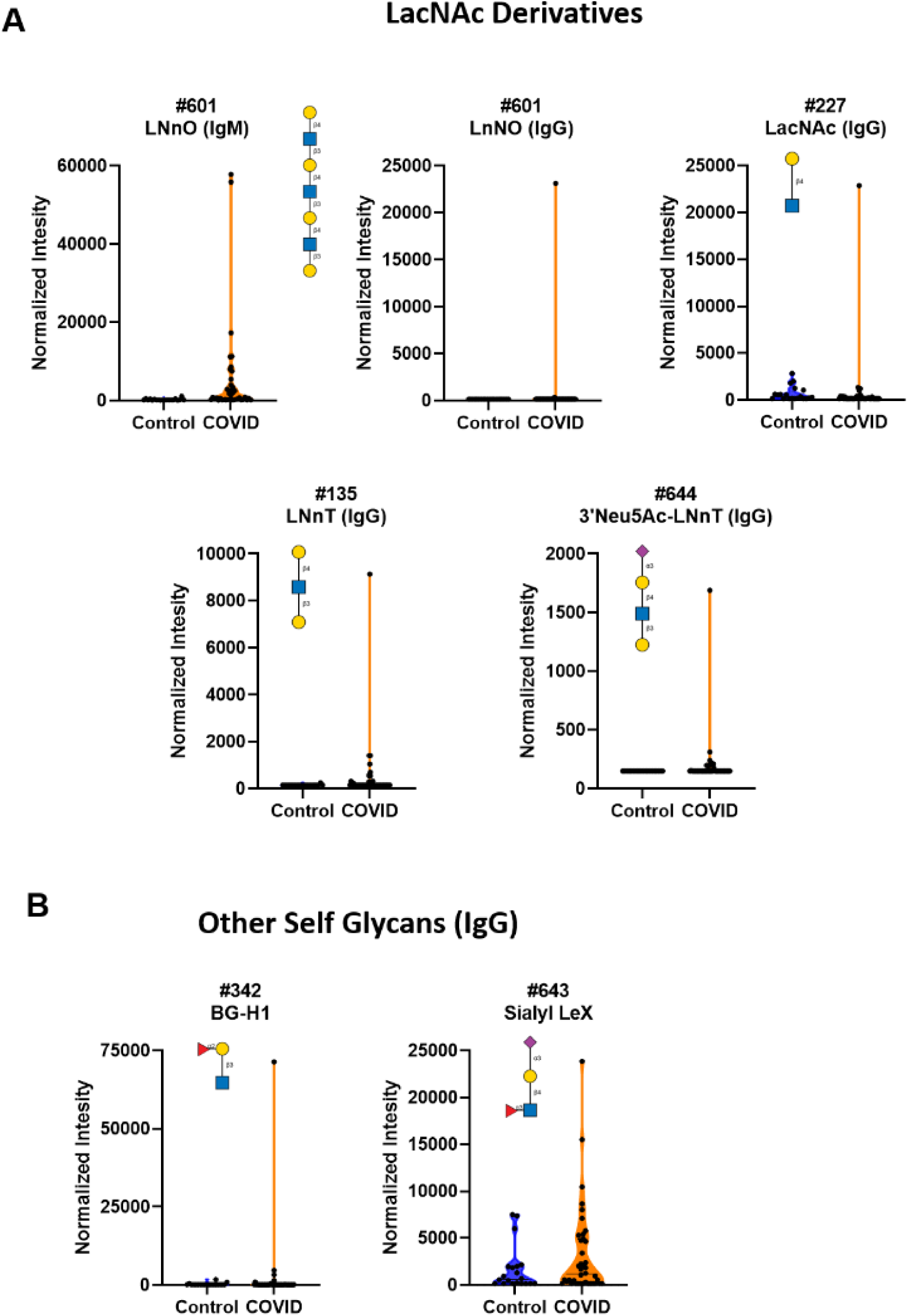
High antibody signals to self-glycans in COVID-19 patient serum. Violin plots show high IgM and IgG signals in COVID-19 patients relative to control donors, with each point representing data from an individual patient: **A)** antibodies to LacNAc derivatives LnNO, LacNAc, LNnT, and sialyl LnNT glycan array components, **B)** antibodies to other self-glycan array components (BG-H1 and Sialyl LeX). See Symbol Key in Figure 1. See Figure 6 for patients with signals to multiple glycans. Glycan structures were created using GlycoGlyph.^59^

Other self-glycans also showed high IgG signals in COVID-19 patients when compared to the control sample set. These glycans included BG-H1 (Fucα1-2Galβ1-3GlcNAcβ) and Sialyl Lewis X (Neu5Acα2-3Galβ1-4[Fucα1-3)GlcNAc) (see Figure 5B and Supplemental Figure S5B).

### Many patients possess antibodies to multiple self-glycans

Most of the gangliosides, *N*-linked glycans, oligomannose glycans, and LNnO discussed in previous sections are found in humans and are considered “self” glycans. To determine if the abnormal antibody signals to these glycans were spread out among the patients or focused in a small subset, we visualized the data in a heat map. Since the signals span a broad range of values, we opted to categorize signals relative to the control group for each glycan component. The signals on the heatmap represent values that are greater than 6 standard deviations above the mean and 10-fold greater than our floor value. As can be seen in the Figure 6, certain patients had high antibodies to multiple types of self-glycans, while others had no antibodies to any of the self-glycans. Of the 7 patients that had high antibodies to at least one *N*-linked glycan, 5 also had high antibodies to one or more gangliosides.

**Figure 6.**
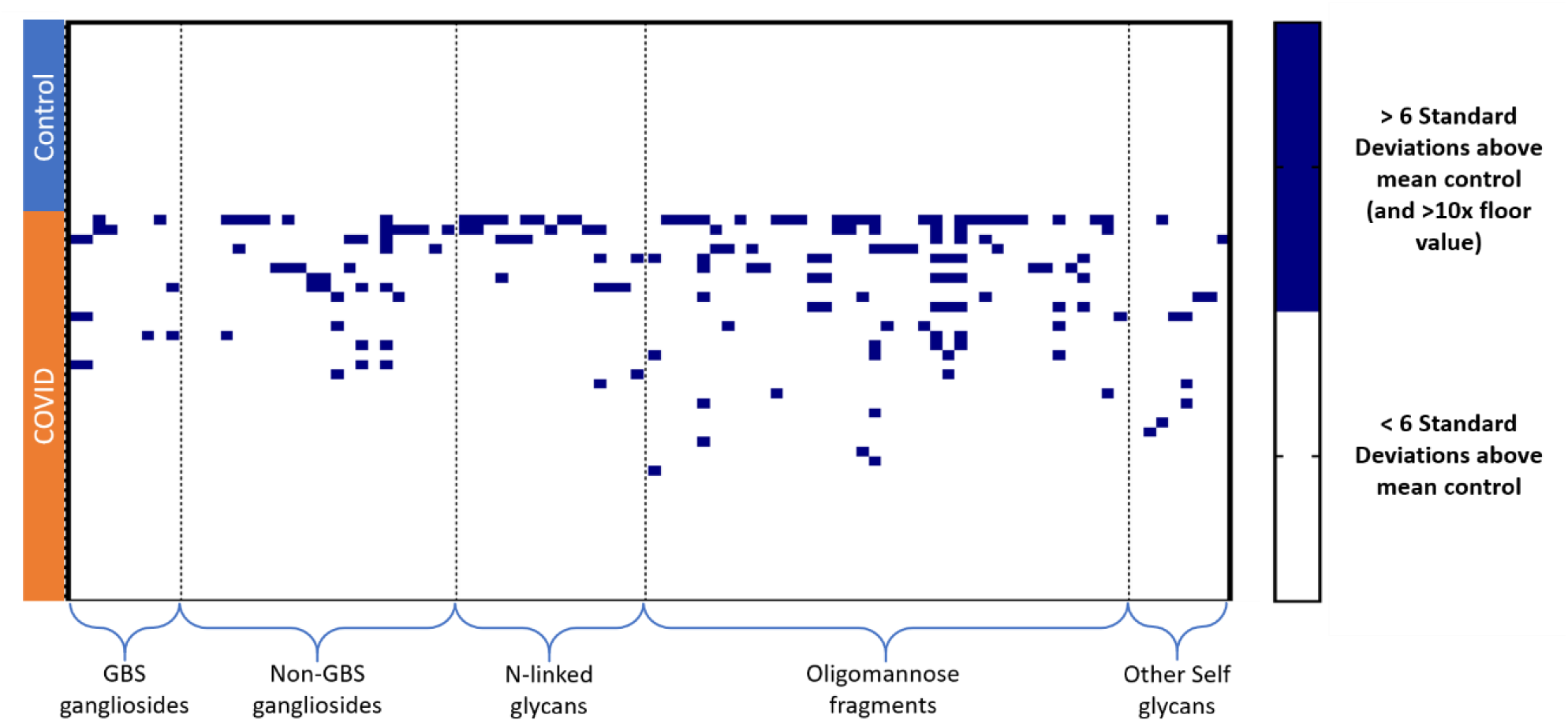
IgG signals from Control and COVID-19 serum samples. Each row represents a patient, each column represents a glycan. Rows are grouped by patient type, columns are grouped by glycan families. Dark blue boxes represent signals that are unusually high (i.e. at least 6 standard deviations above the mean of the control group and at least 10-fold higher than the floor RFU value for our assay). White boxes represent signals that are below that threshold.

### Lower IgG to Sialyl Lewis C, Lewis C, and GN-Lewis C

While there were many glycan families that had high IgG signals among COVID-19 patient samples, there were lower IgG signals to Sialyl Lewis C and Lewis C, and GN-Lewis C glycans (Figure 7 and Figure S6). As can be seen from the violin plots in Figure 7, the lower averages for the COVID-19 cohort were largely driven by a subset of patients with very low signals for these glycans.

**Figure 7.**
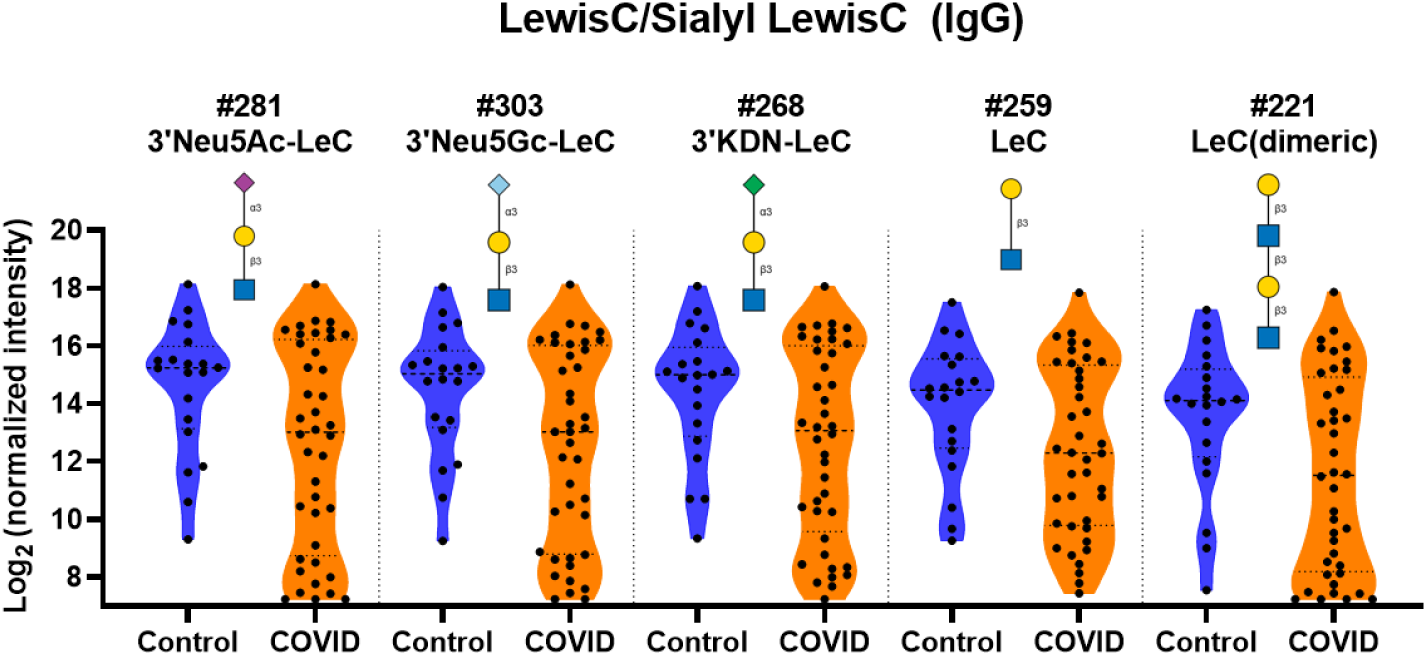
Distribution of IgG signals to Lewis C derivatives. Violin plots show differences in the distribution of IgG signals Lewis C and sialyl Lewis C glycan array components for serum from COVID-19 patients compared to signals seen from serum from control donors, with each point representing data from an individual patient. See Symbol Key in Figure 1. Glycan structures were created using GlycoGlyph.^59^

### Higher IgG but lower IgM to alpha-Gal and other non-human glycans

A previous study by Urra et al. reported an inverse correlation for IgG and IgM antibodies to alpha-Gal[Galα1-3Galβ1-3(4)GlcNAc] and COVID-19 disease severity; those with the most severe outcomes had the lowest levels of α-Gal antibodies.^45^ In their study, COVID-19 patients as a group had lower antibody levels than healthy subjects. Conversely, our results demonstrated higher overall mean α-Gal IgG antibodies (Figure 8 and Figure S7). While we did detect lower IgM antibody signals to α-Gal in COVID-19 samples, this could be due to the overall lower IgM levels seen across almost all glycan antibodies.

**Figure 8.**
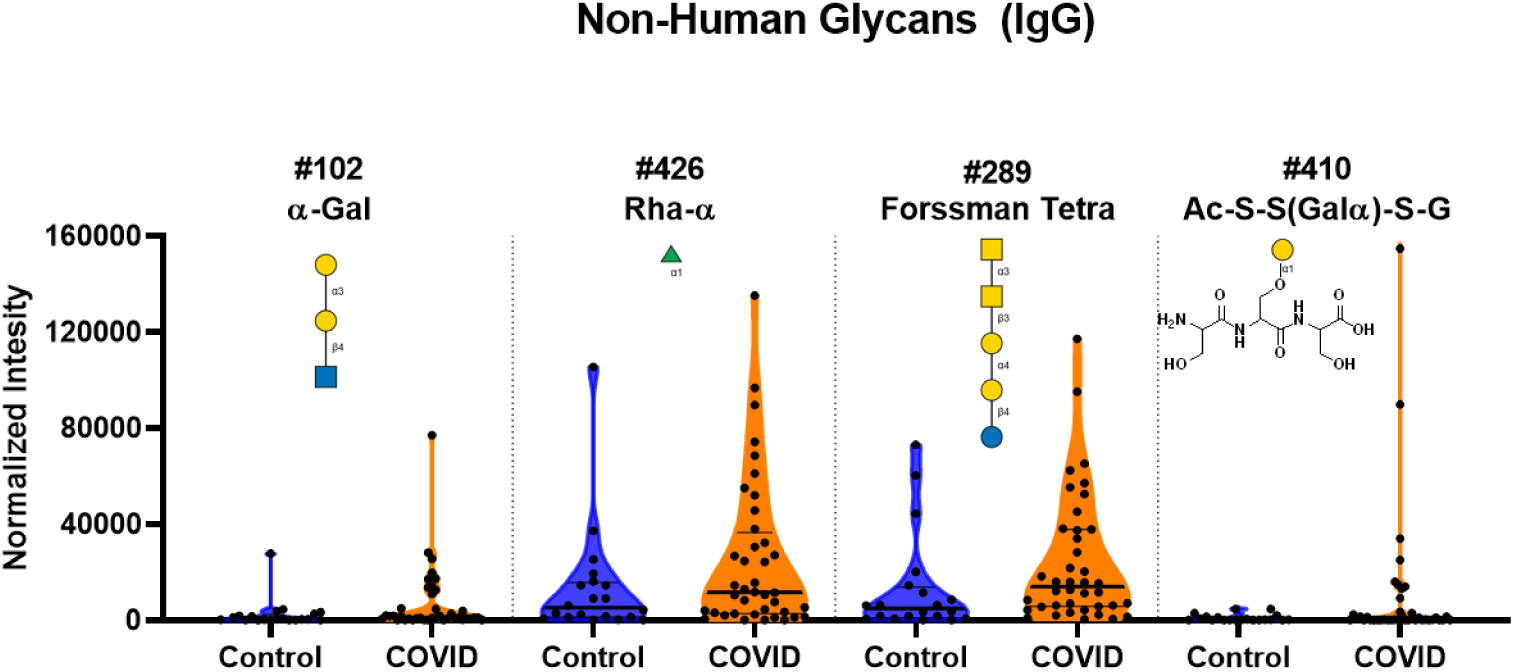
High antibody signals to select non-human glycans in COVID-19 patient serum. Violin plots show high IgG signals to α-Gal, α-rhamnose, Forssman, and Ac-S-S(Galα)-S-G non-human glycan array components for serum from COVID-19 patients compared to baseline signals seen from serum from control donors, with each point representing data from an individual patient. See Symbol Key in Figure 1. Glycan structures were created using GlycoGlyph.^59^

IgG signals to other non-human glycans, such as α-rhamnose, a galactose-modified peptide, and Forssman antigen oligosaccharides, were also higher in COVID-19 patients than controls (Figure 8 and Figure S7). The average signals for COVID-19 patients for the α-rhamnose array components was 1.9-2.5-fold higher than the average signal for the control samples. The average signals for COVID-19 patients were 1.5-3.7-fold higher for the various α-Gal array components compared to the average control signal. The average signal for COVID-19 patents to the Forssman antigens were 1.5-4.4-fold higher than the control samples. In the case of the galactose-modified peptide, nine COVID-19 patients had signals that were unusually high compared to the control samples. No differences in the IgM signals were observed for these glycans.

### IgG and IgM to blood group antigens

There have been several studies that have shown a correlation between blood type and COVID-19 infection rate.^36–39^ In particular, individuals with blood type A have a slightly higher infection rate than those with blood type O. Since serum antibodies to blood group antigens are highly correlated with blood type, we next examined this family of antibodies. Based on reports that blood type A individuals have higher infection rates, we might expect to see lower antibody signals to blood group A antigens. Instead, our results showed higher IgG antibodies to blood group A and B trisaccharide antigens (Figure 9). Other variants of blood group A and B showed similar trends (see Supporting Information, Figure S8). Due to the relatively small nature of our sample size and incomplete patient information about blood type, this higher level of IgG antibodies to blood group A and B could be a random effect.

**Figure 9.**
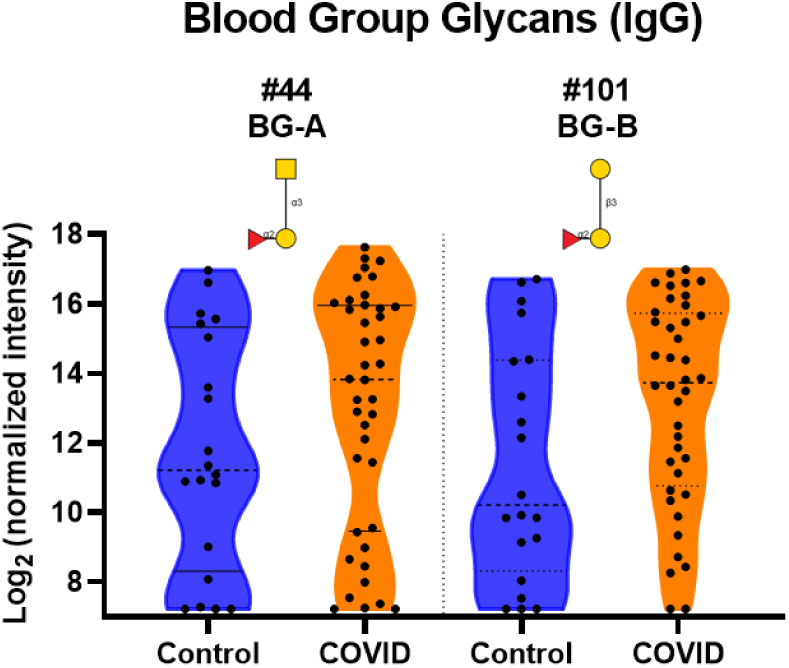
Distribution of IgG signals to Blood Group Antigens. Violin plots show a distribution of higher IgG signals to select Blood Group A and B glycan array components for serum from COVID-19 patients compared to signals seen from serum from control donor, with each point representing data from an individual patient. See Symbol Key in Figure 1. Glycan structures were created using GlycoGlyph.^59^

### Antibodies to *N*-linked glycans bind SARS-CoV-2 spike protein

The results from profiling the serum samples on the glycan array led us to test several mAbs for binding to both subunits and the receptor binding domain (RBD) of the SARS-CoV-2 spike protein. We chose to test antibodies that are known to have binding to oligomannose glycans and A2 since we observed unusually high signals for these antibodies during the array profiling and some monoclonal antibodies to these glycans were available. Several anti-HIV mAbs that bind either oligomannose (PGT126 and PGT128) or A2 (PGT121) fit this category and were tested using an ELISA assay with SARS-CoV-2 spike protein S1, S1 RBD, and S2 subunits on the plate (see Figure 10). PGT128 showed the highest binding to all three spike protein constructs that were tested with best fit apparent *K*D values of 52, 47, and 57 µg/mL (S1, S1 RBD, and S2, respectively). Both PGT126 and PGT121 also bound to all three spike protein constructs, albeit with weaker affinity. Thus, at least some antibodies to *N*-glycans have the potential to recognize glycans as they are presented on the spike protein. None of the antibodies demonstrated neutralization activity (see Supporting Information, Figure S9).

**Figure 10.**
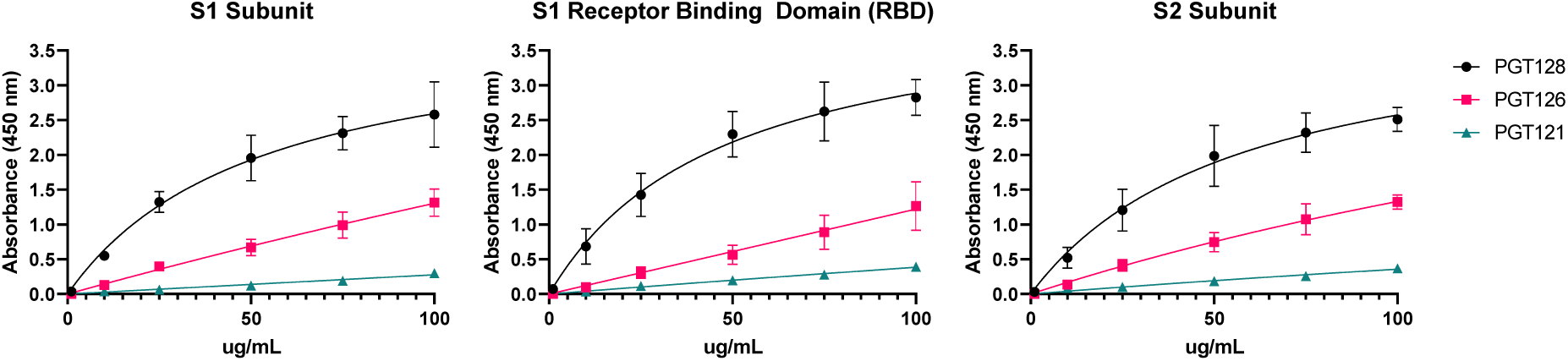
Binding of HIV mAbs to SARS-CoV-2 Spike Protein Fragments. ELISA dilution curve for binding of HIV mAbs PGT128, PGT126, and PGT121 to SARS-CoV-2 spike protein S1 subunit, S1 receptor binding domain, and S2 subunit. Data shown as mean of 2 replicates with error bars showing SEM.

## Discussion

Understanding immune responses to SARS-CoV-2 infection is critical for preventing and treating the disease. For example, SARS-CoV-2 can trigger an overly aggressive immune response leading to excessive damage to the patient, and uncovering this problem has led to the use of the anti-inflammatory agent dexamethasone as an effective treatment for COVID-19.^28^ While there is considerable information being reported on various aspects of the response,^13, 23–25, 43, 44, 61^ such as changes to immune cell populations, cytokine production, and antibodies to proteins, very little is known about immune responses to carbohydrates. Since the surface of the virus is heavily glycosylated,^49–51^ responses to glycans could be triggered, contributing to many aspects of the illness. In addition, pre-existing antibodies to glycans could potentially recognize the virus and influence disease progression. To address these possibilities, we used a large carbohydrate antigen microarray to profile serum anti-glycan IgG and IgM antibody repertoires in COVID-19 patients versus control subjects.

The most distinctive and remarkable differences in COVID-19 patients relative to control subjects were unusually high antibodies to numerous self-carbohydrates, including gangliosides, *N*-linked glycans, LacNAc derivatives (LNnO), blood group H1, and sialyl Lewis X. In many cases, the antibody signals observed in COVID-19 patients were greater than 20 times higher than the largest signal in the control group. In the case of LNnO, the largest COVID-19 patient signal was 154-fold larger than the highest control signal for that glycan. Antibodies to a small subset of gangliosides have been reported previously in several COVID-19 patients.^33, 62, 63^ Our study provides further support of those observations and uncovers antibodies to a much larger assortment of gangliosides/glycolipids than previously reported. In addition to these, we also report many abnormally high antibodies to *N*-linked glycans, LNnO, blood group H1, and sialyl Lewis X, which have not been previously reported in COVID-19 patients. Taken together, our results demonstrate a much more extensive response to self-glycans in a much larger proportion of COVID-19 patients than previously known.

Several lines of evidence indicate that the high anti-glycan antibodies to self-glycans observed in COVID-19 patients are unique and specific to infection. We have investigated anti-glycan antibody repertoires in numerous human serum samples previously, including over 200 healthy subjects and over 100 cancer patients before and after treatment with various cancer vaccines.^56, 64–68^ Based on our prior work, abnormally high antibodies to human gangliosides, *N*-linked glycans, and other self-glycans are uncommon. For example, we did not observed high antibodies to these glycans in ~100 cancer patients prior to or after vaccination with a live-attenuated poxvirus-based vaccine (PROSTVAC-VF),^64, 65^ indicating that they are not due to a general effect of disease or a non-specific effect of viral infection. Some instances where we have observed high antibodies to some of the glycans are HIV infected patients (antibodies to Man9, GT2, and GT3)^66^ and cancer patients immunized with a whole cell cancer vaccine (antibodies to GM2, GM3, Gb5, and sialyl Lewis X).^67^ In these cases, antibodies to self-glycans were present in fewer patients and for fewer glycans than what we observed in COVID-19 patients. In prior studies, we found that serum IgG and IgM levels to nearly all glycans on our array are stable over time frames of up to 3 years,^66, 68^ indicating that high signals in certain patients are not simply due to high variability or random fluctuations over time. Lastly, our prior studies on healthy subjects of varying age indicate that these high antibody populations are not merely due to increasing age.^56^

Antibodies to self-glycans could occur via several possible mechanisms. It is known that antibodies to self-glycans can be induced during certain viral and bacterial infections. For example, autoantibodies to glycans have been reported after infections with *C. jejuni, M. pneumoniae*, *H. influenzae*, cytomegalovirus, Epstein-Barr Virus, Zika virus, chikungunya, and HIV.^69–76^ One mechanism for induction involves molecular mimicry. Certain pathogens produce glycans that are similar to human glycans. Due to the similarity in structure, these glycans can trigger autoantibodies. Another pathway for induction of antibodies to self-glycans occurs when pathogens use host glycosylation machinery to decorate their surface with host glycans. While this process is often used by the viruses to mask themselves from the immune system, response can occur that lead to autoantibodies. A third mechanism for induction of antibodies to self-glycans occurs with enveloped viruses. During the construction and assembly of the viral envelope, host glycoproteins and glycolipids from the endoplasmic reticulum and the Golgi can be incorporated into the envelope along with the viral proteins. When this happens, immune responses to the virus can include antibodies to self-antigens on its surface. Antibodies to glycans in SARS-CoV-2 infected patients could occur via mechanisms two and/or three. Previous studies have shown extensive glycosylation of the spike protein with both *N*-linked and *O*-linked glycans.^49–54^ Therefore, responses to *N*-linked glycans could either be induced by the spike protein or by human glycoproteins incorporated into the envelope. Responses to gangliosides/glycolipids would likely arise via the third mechanism: recognition of glycans incorporated into the SARS-CoV-2 envelope.

Antibodies to self-glycans could be clinically relevant for a variety of reasons. Autoantibodies to self-glycans are associated with a variety of autoimmune disorders.^57, 76–78^ For example, antibodies to gangliosides are often linked to neurological disorders such as Guillain-Barre Syndrome (GBS) and Miller Fisher Syndrome. Gangliosides are expressed at high levels on nerve cells, and antibodies to these glycans can have a variety of effects, including destruction of the neuromuscular junction of nerve cells and disruption of the blood-nerve barrier and/or blood-brain barrier.^79, 80^ Gangliosides also play roles in immune tolerance, signal transduction, and cell adhesion, and antibodies to gangliosides can disrupt these processes as well.^81^ From a clinical perspective, antibodies to GM1, GD1a, GM1b, and GalNAc-GD1a are linked to acute motor axonal neuropathy, and antibodies to GQ1b, GT1a, GD1b, and GD3 are associated with cranial, bulbar, and sensory variants of GBS.^76, 78, 82^ Antibodies to gangliosides are also associated with other diseases such as Alzheimer’s disease, multiple sclerosis, type I diabetes, Crohn’s disease, colitis, and narcolepsy.^76, 78^ Much less is known about clinical effects of antibodies to *N*-linked glycans and other self-glycans, but these glycans are present on numerous cells in the human body and could serve as autoantigens.

Our study has several important implications for treating and preventing COVID-19. One treatment that has recently been granted emergency use authorization is convalescent plasma therapy.^18, 83^ A close variant is anti-coronavirus hyperimmune intravenous immunoglobulin (hIVIG), which has recently entered Phase III clinical trials.^84^ The goal of these approaches is to provide COVID-19 patients with neutralizing antibodies to the virus from patients who have recovered from the disease. Convalescent plasma is typically only screened for a limited set of specific characteristics such as neutralizing antibody titers and the absence of other infectious diseases. While these characteristics are important, our results (and results from others) indicate that screening for potential autoantibodies may be useful to minimize potential complications. For example, one may want to screen plasma for the presence/absence of antibodies to the gangliosides, *N*-linked glycans, and other self-glycans discussed above to ensure that patients receiving convalescent plasma are not being infused with antibodies to these self-glycans.

In addition to treatment, there is also an urgent need to develop a safe and effective SARS-CoV-2 vaccine. Currently, there are numerous vaccines in development using a variety of strategies to initiate immune responses to SARS-CoV-2. The primary measures of success are the reduction of infection rates and the development of neutralizing antibodies. It is possible that some of the vaccines may induce autoantibodies in subsets of patients. Our results, combined with results from other studies,^29–34^ indicate that autoantibodies are a potential complication and that vaccines should be designed to minimize potential autoantibody production. Factors such as the production method and the type of vaccine may be critical. For example, live-attenuated virus or inactivated virus would likely still display a complex assortment of self-glycans to the immune system, providing an opportunity to generate antibodies to self-glycans. Regardless of the type of vaccine, assessing production of potential autoantibodies to glycans, as well as proteins, should be part of the evaluation process.

The results of this study may help to explain some of the unusual symptoms in COVID-19 patients as well as provide insight for developing and choosing treatments. A substantial proportion of COVID-19 patients experience neurological symptoms, such as reduced sense of smell, headaches, muscle pain and spasms as well as delirium, septic encephalopathy, and ischemic stroke.^85, 86^ These symptoms do not appear to be caused by SARS-CoV-2 infection in the brain, as the virus is absent in most cerebrospinal fluid samples.^31^ A variety of other symptoms in COVID-19 patients are not easily explained by direct infection of the affected organ/cells. Many of the symptoms of COVID-19, especially the prolonged symptoms in “long haulers,” resemble autoimmune disorders, and autoantibodies could be key mediators of these symptoms.^31^ In addition to our study focused on antibodies to self-glycans, other studies have shown autoantibodies to a variety of proteins in adult patients with severe COVID-19 symptoms or neurological symptoms.^30, 31^ Understanding the potential roles of autoantibodies may lead to better treatments. For example, Guillain-Barre Syndrome is often treated with intravenous immunoglobulin (IVIG). COVID-19 patients with high antibodies to various gangliosides, and possibly other self-glycans, might also benefit from IVIG. Additional studies will be needed to evaluate this hypothesis.

Our study further illustrates how patient symptoms and immune responses to SARS-CoV-2 infection can vary widely. While some patients have neurological complications and other symptoms associated with autoantibodies, others have much milder symptoms. In our study, patient symptoms were unknown, but some patients had broad responses to self-glycans while others had none. For example, five of the COVID-19 patients accounted for over half of all the unusually high signals for antibodies to self-glycans. This result is consistent with a model wherein tolerance is broken in certain patients, leading to widespread production of autoantibodies to an assortment of self-antigens.

Beyond the self-glycans, we observed substantial differences between COVID-19 patients and control subjects for a variety of other glycans, including Lewis C/Sialyl Lewis C, rhamnose, the Forssman antigen, and a glycopeptide with galactose α-linked to a serine residue. It is not yet clear why there would be differences in antibodies to these glycans. Secondary infections are a possibility, but more studies will be needed to better understand the basis of these differences.

Several limitations of this study should be mentioned. First, our glycan microarray only contains a small portion of the glycans found in the human glycome. Thus, there may be other important anti-glycan antibody populations that were not detected. Second, our study included a relatively small cohort of 40 COVID-19 patients and 20 healthy controls. In other work, we have profiled serum anti-glycan antibodies in hundreds of healthy subjects, so our understanding of normal antibody repertoires draws from considerable experience.^56, 87^ In contrast, these are the first 40 COVID-19 patients we have evaluated, and additional testing will be helpful to more fully investigate the findings in this study. Third, information about patient symptoms and outcome were not available. Consequently, follow up studies will be needed to evaluate potential correlations between symptoms and anti-glycan antibody repertoires. Additional studies to address these limitations are currently underway.

Lastly, our study highlights the importance of studying immune responses to carbohydrates. Glycans are one of the major families of antigens found on SARS-CoV-2 and other viruses, but responses to these antigens are often difficult to study. By profiling serum antibodies with a large and diverse carbohydrate antigen microarray, we were able to rapidly identify abnormally high antibodies to a variety of self-glycans. These results provide new insight into the immune response to SARS-CoV-2 and illustrate the importance of studying antibodies to host antigens in addition to viral antigens. The results also highlight key factors/concerns for developing vaccines and treatments for COVID-19 and provide a more complete understanding of the risks associated with SARS-CoV-2 infection, which is critical for making informed health decisions.

## Materials and Methods

### Serum Samples

Publicly available, de-identified serum samples from 40 individuals with SARS-CoV-2 infections and 10 healthy donors were purchased from RayBiotech, Inc. (Peachtree Corners, GA). These samples were collected on-site at multiple RayBiotech locations within the US. All patients designated to be infected with SARS-CoV-2 were symptomatic. Ten additional healthy donor serum samples were obtained from Valley Biomedical Products and Services (Winchester, VA). All non-COVID-19 samples were collected prior to the SARS-CoV-2 pandemic. Among the COVID-19 positive samples, 10 were IgM positive, 10 were IgG positive and 20 were not specified as either IgM or IgG positive. The reference serum was pooled from 10 samples purchased from Valley Biomedical Products and Services. Samples were stored at −70°C prior to use.

### Microarray fabrication and assay

The glycan microarrays were fabricated as previously described.^88, 89^ The microarray contained 816 array components and included a variety of human glycans (*N-*linked glycans, *O*-linked glycans, and glycan portions of glycolipids), non-human glycans, glycopeptides, and glycoproteins. Each array component was printed in duplicate to produce a full array, and 8 copies of the full array were printed on each slide. Prior each experiment, each microarray slide was scanned in an InnoScan 1100 AL fluorescence scanner to check for any defects and missing print spots. The slides were fitted with an 8-well module (Sigma-Aldrich) to allow 8 independent assays on each slide. In the assay, arrays were blocked with 3% BSA in PBS buffer (400 µL/well) overnight at 4 °C, then washed six times with PBST buffer (PBS with 0.05% v/v Tween 20). Serum samples diluted at 1:50 in 3% BSA and 1% HSA in PBST were added onto each slide (100 µL/well). To minimize technical variations, all samples were assayed in duplicate on separate slides. After agitation at 100 rpm for 4 hours at 37 °C, slides were washed six times with PBST (200 µL/well). The bound serum antibodies were detected by incubating with Cy3 anti-Human IgG and DyLight 647 anti-human IgM (Jackson ImmunoResearch) at 3 µg/mL in PBS buffer with 3% BSA and 1% HSA (100 µL/well) under agitation at 37 °C for 2 hours. Slides were covered with aluminum foil to prevent photobleaching. After washing with PBST eight times (200 µL/well), the slides were removed from the modules and soaked in PBST for 5 min prior to being dried by centrifugation at 1000 rpm (112 × *g*) for 10 minutes. Slides were then scanned with an InnoScan 1100 AL (Innopsys) at 5 µm resolution. The photomultiplier tube (PMT) settings were the same for all experiments to limit unintentional signal variation. Slides were scanned at “high” and “low” PMT settings (for the 532 nm laser, high pmt = 5 and low = 1; for the 635 nm laser, high = 25 and low = 9) to increase the dynamic range and appropriately scale-saturated components. The fluorescence intensity of each array spot was quantified with GenePix Pro 7 software (Molecular Devices). Any features marked as missing or defective in the prescan were excluded from further analysis. The local background corrected median was used for data analysis, and spots with intensity lower than 150 RFU (1/2 the typical IgM background) were set to 150. The signals for replicate spots on duplicate wells were averaged and log-transformed (base 2) for future analysis. Full microarray data can be found in the Supporting Excel file.

### ELISA Assay

Recombinant SARS-CoV-2 S1 subunit protein (RBD) (Raybiotech, Inc.), recombinant SARS-CoV-2 S1 subunit protein (full length) (RayBiotech, Inc.), or recombinant SARS-CoV-2 S2 subunit protein (full length) (RayBiotech, Inc.) were plated at 5 µg/mL in PBS buffer, pH 7.4 into the desired wells of a 96-well clear, flat-bottomed ELISA plate (ThermoFisher Scientific, Nunc MaxiSorp^™^). The plates were covered with an adhesive foil and stored at 4°C overnight. The plates were emptied and washed four times with PBS buffer (200 µL/well). The plates were blocked with 3% BSA in PBS buffer (100 µL/well) for 2 hours at room temperature. The plates were washed four times with PBS buffer (200 µL/well). Each of the antibody stocks (PGT121, PGT126, PGT128, NIH AIDS Reagent Program) were diluted to 100 µg/mL diluted 5-fold in 1% BSA in PBS buffer. Antibody solutions were added to the plate (50 µL/well) and incubated at 37°C for 2 hours with gentle agitation. The plates were emptied and washed six times with PBS buffer (200 µL/well). A solution of peroxidase affinity pure goat anti-human IgG (1:2500 dilution, JacksonImmuno) in 1% BSA in PBS buffer was added to each well (50 µL/well) and incubated at 37°C for 2 hours with gentle agitation. The plates were emptied and washed six times with PBS buffer (200 µL/well). After plates were washed, TMB ELISA substrate (50 µL/well, high sensitivity, abCam) was added to each well and allowed to sit at room temperature for 30 minutes before a stop solution (50 µL/well, 2M H2SO4) was added to each well. The absorbance at 450 nm was read using a BioTek Biosynergy 2.

## Supporting information

Full array data

## Acknowledgements

We thank the Consortium for Functional Glycomics (GM62116; The Scripps Research Institute), X. Huang (Michigan State University), T. Tolbert (University of Kansas), Lai-Xi Wang (University of Maryland), J. Barchi (National Cancer Institute), T. Lowary (University of Alberta), Beat Ernst (University of Basel), Omicron Biochemicals Inc., GlycoHub, and Glycan Therapeutics for generously contributing glycans for the array. The following reagents were obtained through the NIH AIDS Reagent Program, Division of AIDS, NIAID, NIH: Anti-HIV-1 gp120 Monoclonal (PGT121) from IAVI (cat# 12343); Anti-HIV-1 gp120 Monoclonal (PGT126) from IAVI (cat# 12344); Anti-HIV-1 gp120 Monoclonal (PGT128) from IAVI.^90, 91^ This work was supported by the Intramural Research Program of the National Cancer Institute, NIH.

