## Supplementary Methods and Figures for "Abnormal antibodies to self-carbohydrates in SARS-CoV-2 infected patients"

**for**

**Supplementary Materials and Methods**

*Measuring Total IgM*

Invitrogen IgM Human Uncoated ELISA Kit (88-50620) was purchased from Life Technologies and used according to the user manual. Briefly, flat-bottomed ELISA pate (ThermoFisher Scientific, Nunc MaxiSorp™) were coated with capture antibody in Coating Buffer (1:250 dilution, 100 µL /well). The plates were covered in adhesive foil and stored at 4°C overnight. The plates were emptied and washed twice with Wash Buffer (400 µL /well). The plates were blocked with Blocking Buffer (200 µL /well) for 2 hours at room temperature. The plates were emptied and washed twice with Wash Buffer (400 µL /well). The human IgM standard was reconstituted and added to the plate in 2-fold serial dilutions (in Assay Buffer A) in duplicate. The serum samples were added to the plate (1:20,000 dilution in Assay Buffer A). The plates were covered in adhesive foil and incubated at room temperature for 2 hours on a shaker at 200 rpm. The plates were emptied and washed four times with Wash Buffer (400 µL /well). The detection antibody (1:250 in Assay Buffer A, 100 µL /well) was added to the plate. The plates were covered in adhesive foil and incubated at room temperature for 1 hour on a shaker at 200 rpm. The plates were emptied and washed four times with Wash Buffer (400 µL /well). Substrate Solution (100 µL /well) was added to the plate and incubated for 15 minutes then the Stop Solution (100 µL /well) was added to each well. The pate was read at 450 nm using a BioTek Biosynergy 2, and a standard curve was created from the standard samples and used to calculate the concentrations of IgM in the serum samples.

*Neutralization Assay*

COVID-19 Spike-ACE2 binding assay kit (CoV-SACE2) was purchased from RayBiotech, Inc. (Peachtree Corners, GA) and used according to user manual. Briefly, stock solutions were made according to instructions and sample solutions containing ACE2 at a single concentration and each antibody (PGT121, PGT126, PGT128) at two concentrations (20 µg/mL, 50 µg/mL) were made. Sample solutions (50 µL/well) were added to the plate and incubated at room temperature for 2.5 hours with shaking. The plates were emptied and washed four times with 1x Wash Solution (200 µL/well). The detection antibody (anti-ACE2, 100 µL/well) was added to the plate and incubated at room temperature for 1 hour with shaking. The plates were emptied and washed four times with 1x Wash Solution (200 µL/well). HRP-conjugated anti-goat IgG (1:1000 dilution, 100 µL/well) was added to the plate and incubated at room temperature for 1 hour with shaking. The plates were emptied and washed four times with 1x Wash Solution (200 µL/well). TMB One-Step Substrate Reagent (100 µL/well) was added to each well and incubated at room temperature for 30 minutes in the dark with shaking. Stop Solution (50 µL/well) was added to each well. The absorbance at 450 nm was read using a BioTek Biosynergy 2.

**Supplementary Figures**

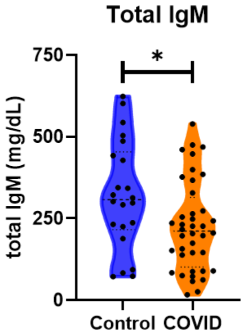


**Figure S1. Total IgM from serum samples.** Violin plots of the total measured IgM from control and COVID serum samples. Mean IgM of control samples was 332.0 mg/dl. Mean of IgM of COVID samples was 226.3 mg/dl. Unpaired t-test with Welch's correction. *, p=0.0222.

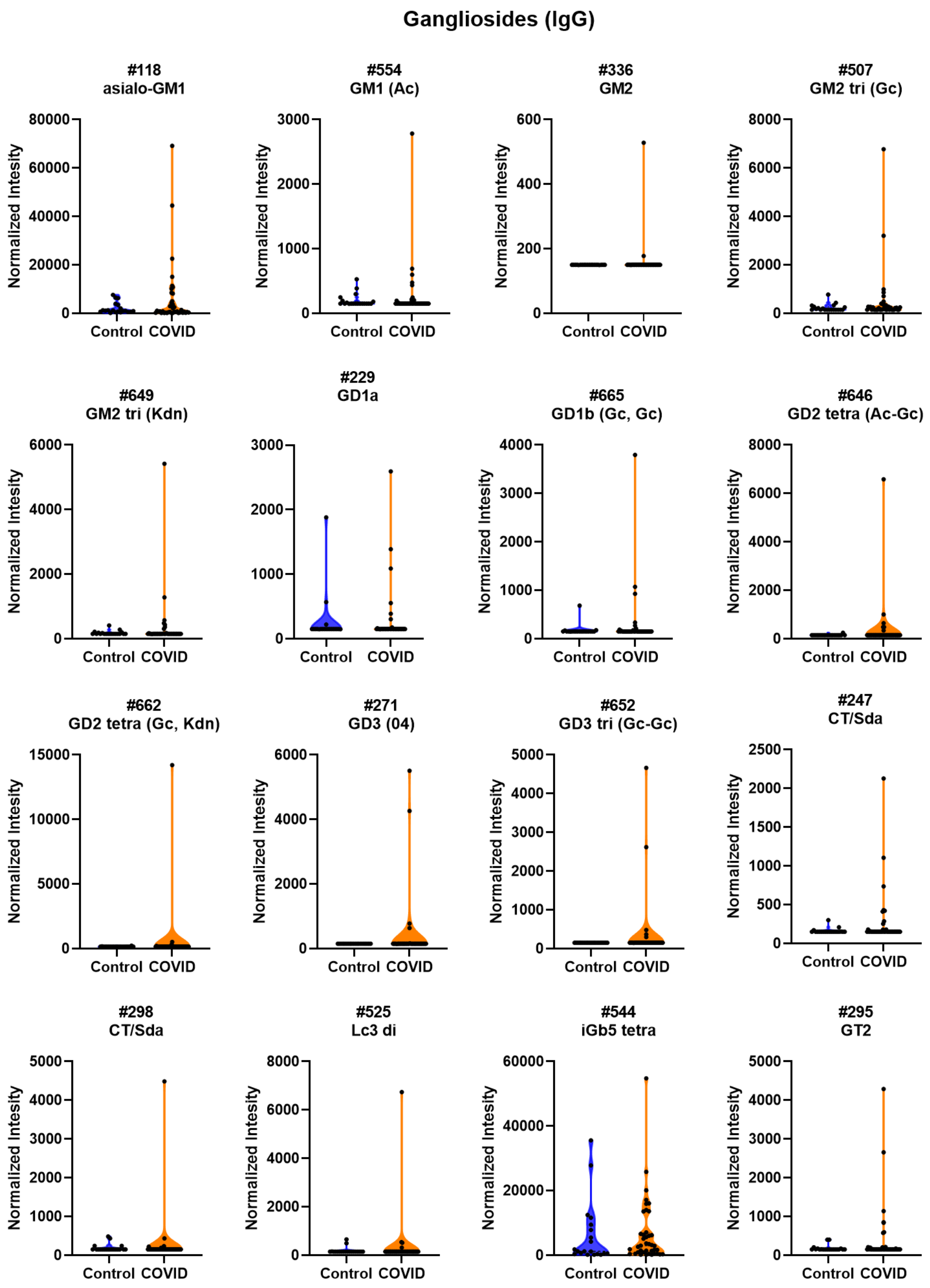


**Figure S2. High antibody signals to select ganglioside glycans in COVID patient serum.** Violin plots show several high IgG signals to ganglioside glycan array components for serum from COVID patients compared to baseline signals seen from serum from control donors. Some components have the same name but differ in either the linker or the density of glycan moieties on the surface or both. Asialo-GM1: Galβ1-3GalNAcβ1-4Galβ; GM1 (Ac): Galβ1-3GalNAcβ1-4(Neu5Acα2-3)Galβ; GM2: Galβ1-3GalNAcβ1-4(Neu5Acα2-3)Galβ; GM2 tri (Gc): Neu5Gcα2-3(GalNAcβ1-4)Galβ; GM2 tri (Kdn): Kdnα2-3(GlcNAcβ1-4)Galb1; GD1a: Neu5Acα2-3[Neu5Acα2-3Galβ1-3GalNAcβ1-4]Galβ1-4Glcβ; GD1b (Gc,Gc): Neu5Gcα2-8Neu5Gcα2-3(Galβ1-3GlcNAcβ1-4)Galβ; GD2 tetra (Ac-Gc): Neu5Acα2-8Neu5Gcα2-3(GlcNAcβ1-4)Galβ; GD2 tetra (Gc, Kdn): Neu5Acα2-8Kdnα2-3(GlcNAcβ1-4)Galβ; GD3: Neu5Acα2-8Neu5Acα2-3Galβ1-4Glcβ; GD3 tri (Gc-Gc): Neu5Gcα2-8Neu5Gcα2-3Galβ; CT/Sda: Neu5Acα2-3(GalNAcβ1-4)Galβ1-4GlcNAcβ; Lc3 di: GlcNAcβ1-3Galβ; iGb5 tetra: Galβ1-3GalNAcβ1-3Galα1-3Galβ; GT2: Neu5Acα2-8Neu5Acα2-8Neu5Acα2-3[GalNAcβ1-4]Galβ1-4Glcβ

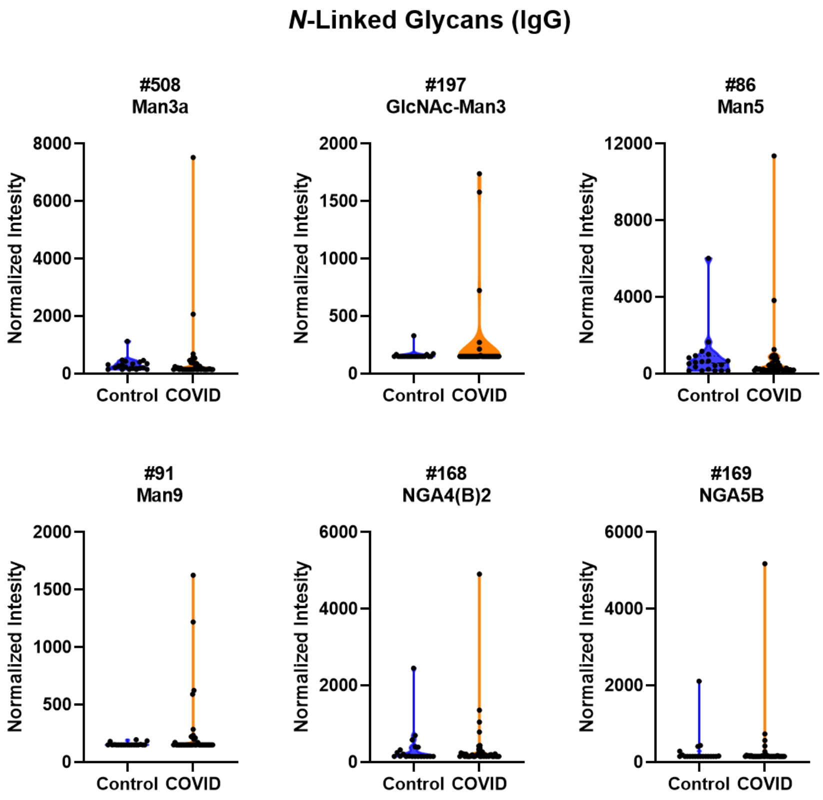


**Figure S3. High IgG signals to select *N*-Linked Glycans in COVID patient serum.** Violin plots show several high IgG signals to select *N*-linked glycan array components for serum from COVID compared to baseline signals seen from serum from control donors. Some components have the same name but differ in either the linker or the density of glycan moieties on the surface or both. Man3a: Mana1-2Mana1-3Manβ1-4GlcNAcβ; GlcNAc-Man3: Manα1-6(GlcNAcβ1-2Manα1-3)Manβ1-4GlcNAcβ; Man5: Manα1-6(Manα1-3)Manα1-6(Manα1-3)Manβ1-4GlcNAcβ; Man9: Manα1-2Manα1-6(Manα1-2Manα1-3)Manα1-6(Manα1-2Manα1-2Manα1-3)Manβ1-4GlcNAcβ; NGA(B)2: GlcNAcβ1-2(GlcNAcβ1-4)(GlcNAcβ1-6)Manα1-6[GlcNAcβ1-2Manα1-3](GlcNAcβ1-4)Manβ1-4GlcNAcβ; NGA5B; GlcNAcβ1-2(GlcNAcβ1-4)(GlcNAcβ1-6)Manα1-6[GlcNAcβ1-2(GlcNAcβ1-4)Manα1-3](GlcNAcβ1-4)Manβ1-4GlcNAcβ.

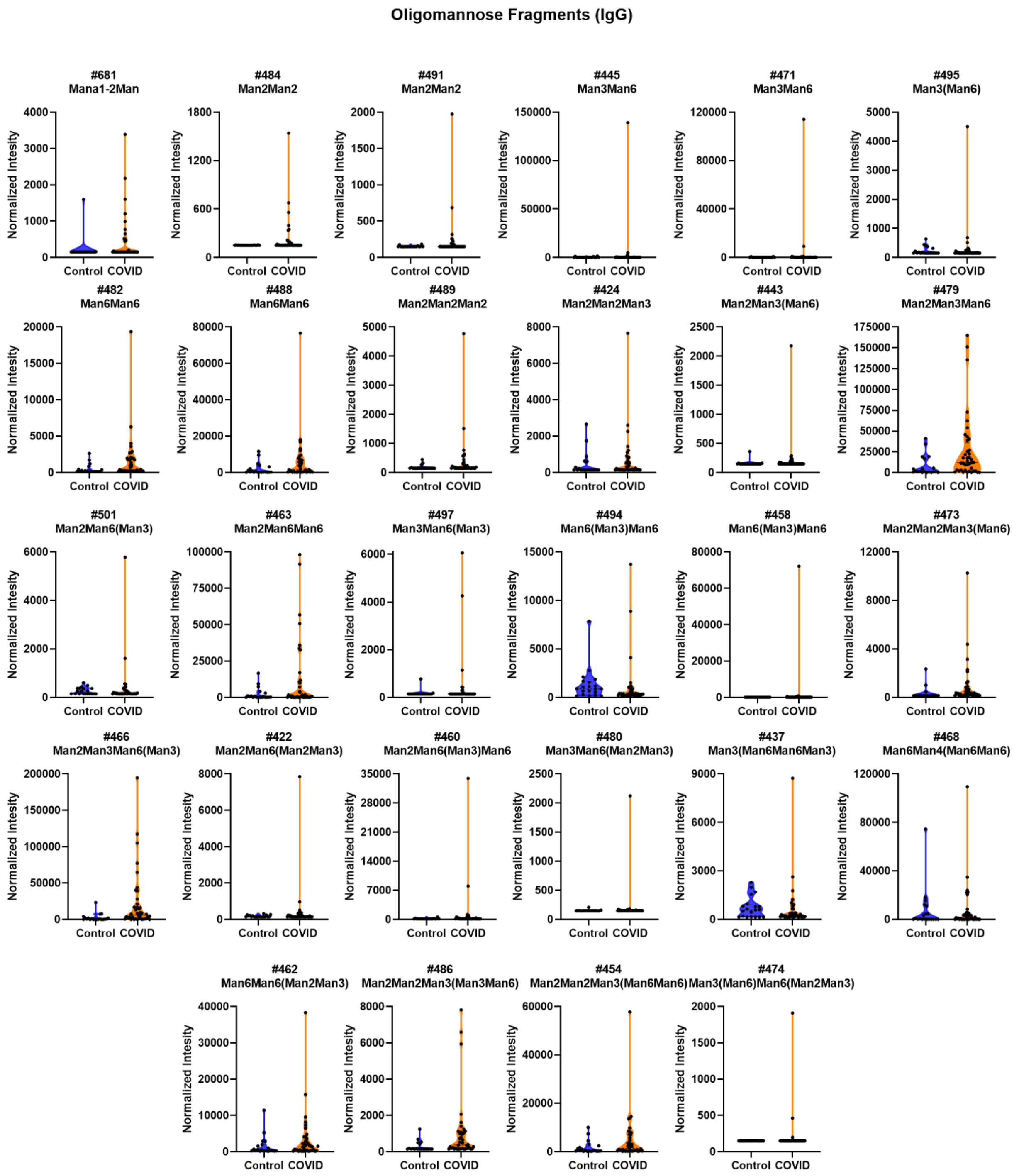


**Figure S4. High IgG signals to select oligomannose fragments in COVID patient serum.** Violin plots show several high IgG signals to select oligomannose glycan array components for serum from COVID compared to baseline signals seen from serum from control donors. Some components have the same name but differ in either the linker or the density of glycan moieties on the surface or both. Mana1-2Man: Manα1-2Man; Man2Man2: αMan(1-2)αMan(1-2); Man3Man6: αMan(1-3) αMan(1-6); Man3(Man6): αMan(1-3)[αMan(1-6)]; Man6Man6: αMan(1-6)αMan(1-6); Man2Man2Man2: αMan(1-2)αMan(1-2)αMan(1-2); Man2Man2Man3: αMan(1-2)αMan(1-2)αMan(1-3); Man2Man3(Man6): αMan(1-2)αMan(1-3)[αMan(1-6)]; Man2Man3Man6: αMan(1-2)αMan(1-3)αMan(1-6);

Man2Man6(Man3): αMan(1-2)αMan(1-6)[αMan(1-3)]; Man2Man6Man6: αMan(1-2)αMan(1-6)αMan(1-6); Man3Man6(Man3): αMan(1-3)αMan(1-6)[αMan(1-3)]; Man6(Man3)Man6): αMan(1-6)[αMan(1-3)]αMan(1-6); Man2Man2Man39Man6): αMan(1-2)αMan(1-2)αMan(1-3)[αMan(1-6)]; Man2Man3Man6(Man3): αMan(1-2)αMan(1-3)αMan(1-6)[αMan(1-3)]; Man2Man6(Man2Man3): αMan(1-2)αMan(1-6)[αMan(1-2)αMan(1-3)]; Man2Man6(Man3)Man6: αMan(1-2)αMan(1-6)[αMan(1-3)]αMan(1-6); Man3Man6(Man2Man3): αMan(1-3)αMan(1-6)[αMan(1-2)αMan(1-3)]; Man3(Man6Man6Man3): αMan(1-3)[αMan(1-6)]αMan(1-6)[αMan(1-3)]; Man6Man4(Man6Man6): αMan(1-6)αMan(1-4)[αMan(1-6)αMan(1-6)]; Man6Man6(Man2Man3): αMan(1-6)αMan(1-6)[αMan(1-2)αMan(1-3)]; Man2Man2Man3(Man3Man6): αMan(1-2)αMan(1-2)αMan(1-3)[αMan(1-3)αMan(1-6)]; Man2Man2Man3(Man6Man6): αMan(1-2)αMan(1-2)αMan(1-3)[αMan(1-6)αMan(1-6)]; Man3(Man6)Man6(Man2Man3): αMan(1-3)[αMan(1-6)]αMan(1-6)[αMan(1-2)αMan(1-3)]

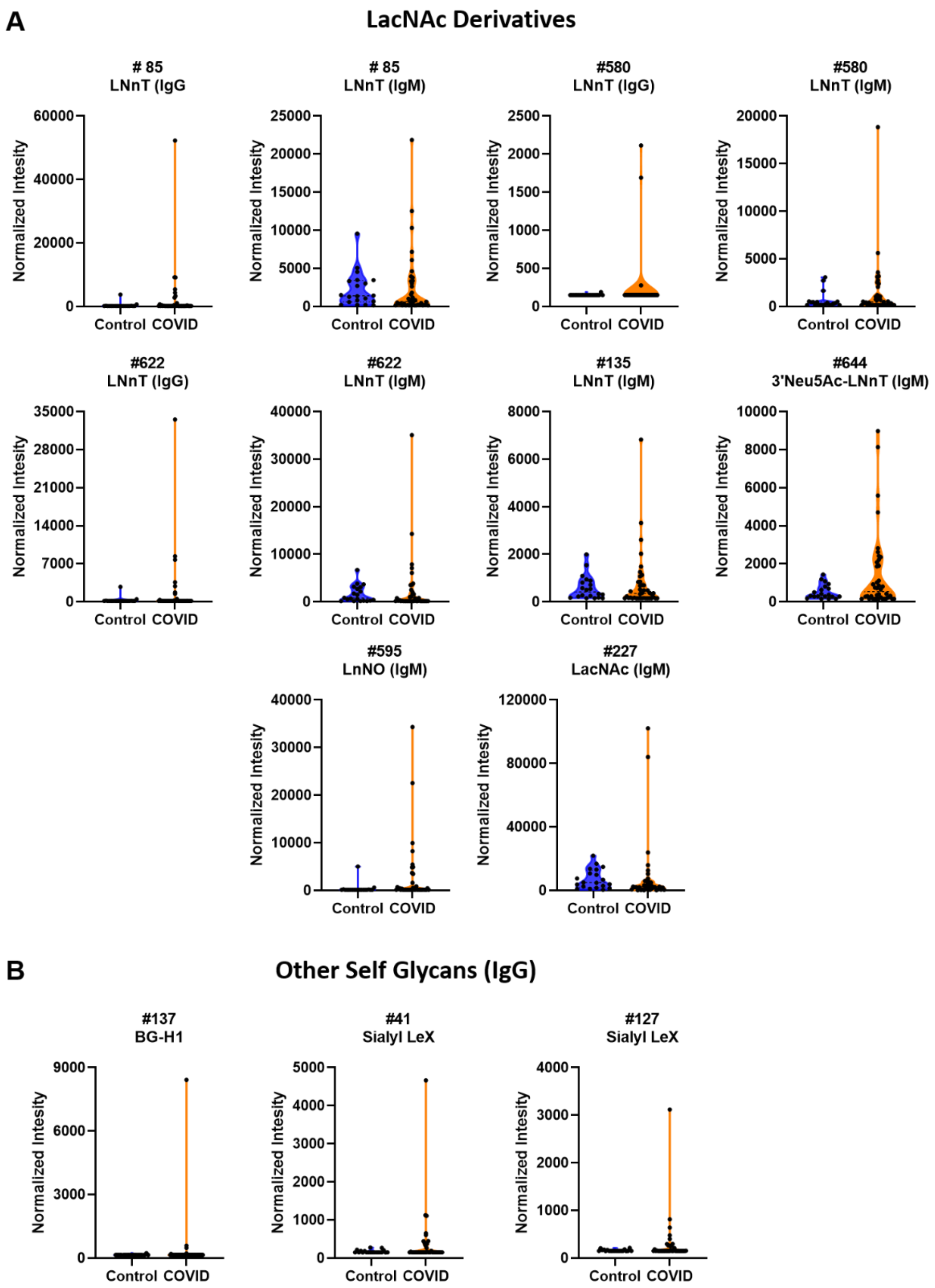


**Figure S5. High antibody signals to self glycans in COVID patient serum.** These glycans are similar to the ones in the main paper but differ in the linker and/or the density of glycan moieties on the surface. **A)** Violin plots show several high IgG and IgM signals to glycan array components that are LacNAc derivatives for serum from COVID compared to baseline signals seen from serum from control donors. **B)** Violin plots show several high IgG signals to self glycan array components (BG-H1 and Sialyl LeX) for serum from COVID compared to baseline signals seen from serum from control donors. LNnT: Galβ1-4GlcNAcβ1-3Galβ; 3'Neu5Ac-LNnT: Neu5Acα2-3Galβ1-4GlcNAcβ1-3Galβ; LNnO: Galβ1-4GlcNAcβ1-3Galβ1-4GlcNAcβ1-3Galβ1-4GlcNAcβ1-3Galβ; LacNAc: Galβ1-4GlcNAcβ; BG-H1: Fucα1-2Galβ1-3GlcNAcβ1; Sialyl LeX: Neu5Acα2-3Galβ1-4[Fucα1-3)GlcNAcβ

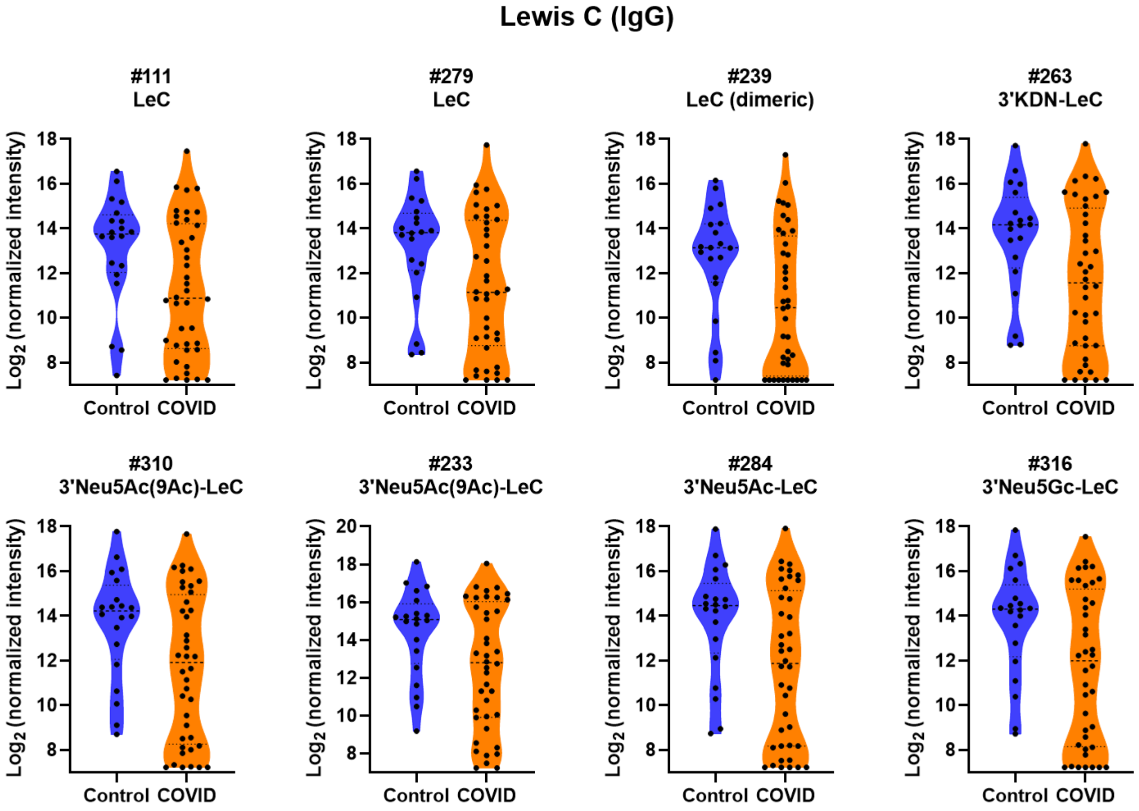


**Figure S6. Distribution of IgG signals to Lewis C derivatives.** Violin plots show differences in the distribution of IgG signals Lewis C and sialyl Lewis C glycan array components for serum from COVID patients compared to signals seen from serum from control donors. Some components have the same name but differ in either the linker or the density of glycan moieties on the surface or both. LeC: Galβ1-3GlcNAcβ; LeC (dimeric): Galβ1-3GlcNAcβ1-3Galβ1-3GlcNAcβ; 3'KDN-LeC: Kdnα2-3Galβ1-3GlcNAcβ; 3'Neu5Ac(9Ac)-LeC: Neu5Ac(9Ac)α2-3Galβ1-3GlcNAcβ; 3'Neu5Ac-LeC: Neu5Acα2-3Galβ1-3GlcNAcβ; 3'Neu5Gc-LeC: Neu5Gcα2-3Galβ1-3GlcNAcβ

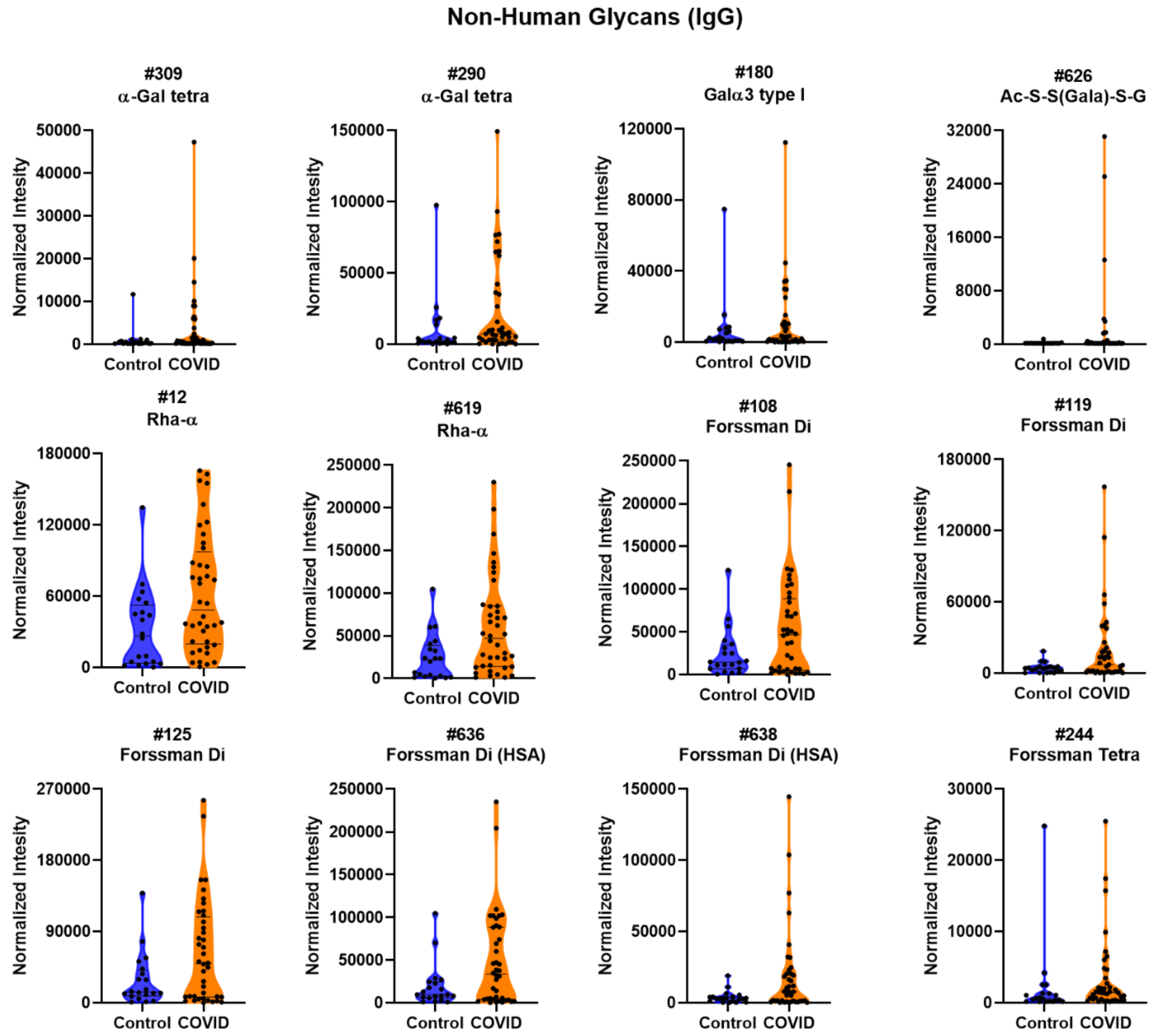


**FigureS7. High antibody signals to select non-human glycans in COVID patient serum.** Violin plots show high IgG signals to α-Gal, α-rhamnose, Forssman, and Ac-S-S(Galα)-S-G non-human glycan array components for serum from COVID compared to baseline signals seen from serum from control donors. Some components have the same name but differ in either the linker or the density of glycan moieties on the surface or both. α-Gal tetra: Galα1-3Galb1-4GlcNAcβ1-3Galβ; Galα3 type I: Galα1-3Galβ1-3GlcNAcβ; Ac-S-S(Gala)-S-G: Serine-Serine$^{(\alpha Gal)}$-Serine; Rha-α: Rha-α; Forssman Di: GalNAcα1-3GalNAcβ; Forssman Tetra: GalNAcα1-3GalNAcβ1-3Gala1-4Galβ

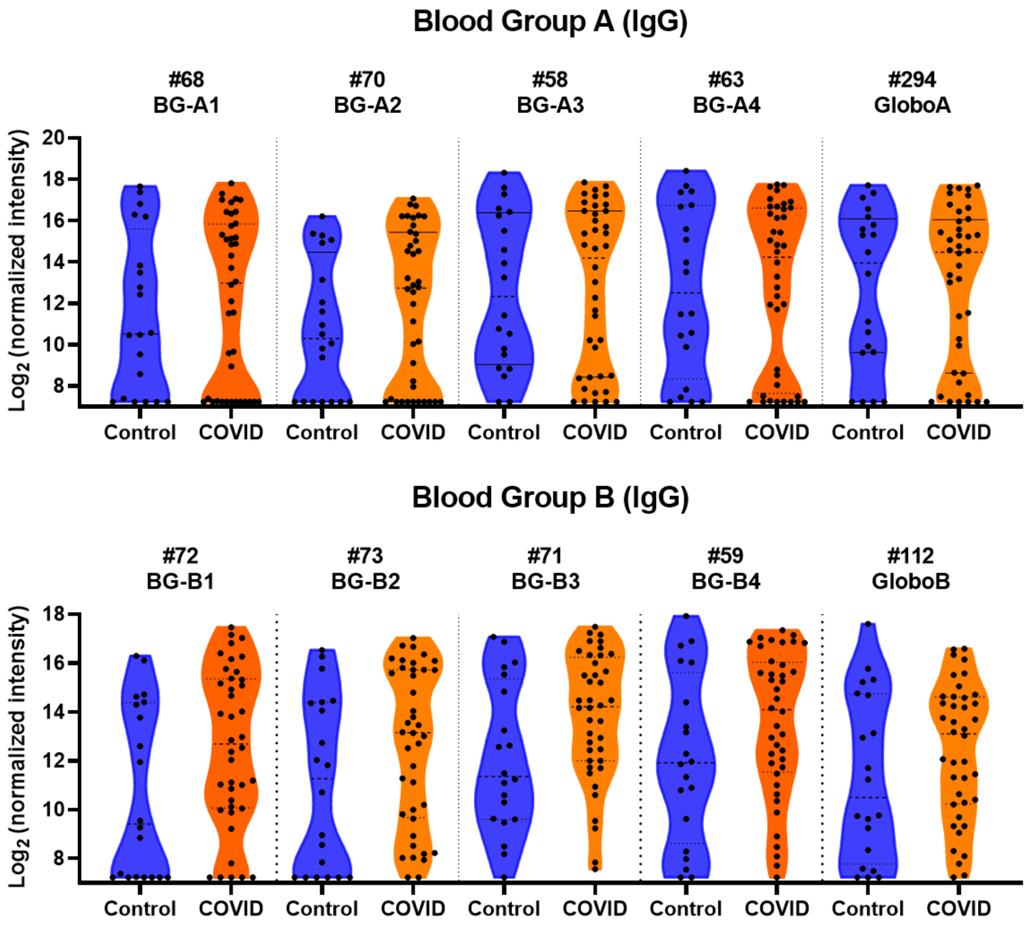


**Figure S8. Distribution of IgG signals to Blood Group Antigens.** Violin plots show a distribution of higher IgG signals to select Blood Group A and B glycan array components for serum from COVID patients compared to signals seen from serum from control donor. BG-A1: GalNAcα1-3(Fucα1-2)Galβ1-3GlcNAcβ; BG-A2: GalNAcα1-3(Fucα1-2)Galβ1-4GlcNAcβ; BG-A3: GalNAcα1-3(Fucα1-2)Galβ1-3GalNAcα; BG-A4: GalNAcα1-3(Fucα1-2)Galβ1-3GalNAcβ; GloboA: GalNAcα1-3(Fucα1-2)Galβ1-3GalNAcβ1-3Galα1-4Galβ1; BG-B1: Galα1-3(Fucα1-2)Galβ1-3GlcNAcβ; BG-B2: Galα1-3(Fucα1-2)Galβ1-4GlcNAcβ; BG-B3: Galα1-3(Fucα1-2)Galβ1-3GalNAcα; BG-B4: Galα1-3(Fucα1-2)Galβ1-3GalNAcβ; GloboB: Galα1-3(Fucα1-2)Galβ1-3GalNAcβ1-3Galα1-4Galβ

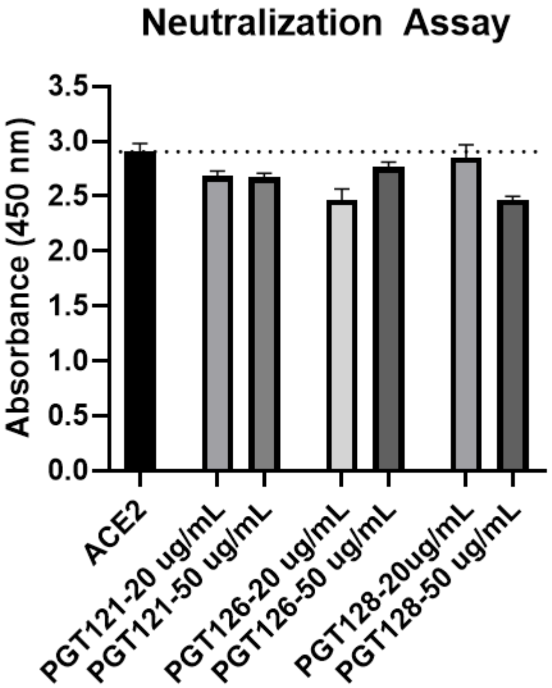


**Figure S9. Neutralization of HIV mAbs against SARS-CoV-2 Spike Protein RBD.** An ELISA assay was used to measure the ability of several HIV mAbs to inhibit the binding of the SARS-CoV-2 Spike Protein RBD to the human ACE2 receptor.
